# Scalable unified framework of total and allele-specific counts for cis-QTL, fine-mapping, and prediction

**DOI:** 10.1101/2020.04.22.050666

**Authors:** Yanyu Liang, François Aguet, Alvaro Barbeira, Kristin Ardlie, Hae Kyung Im

## Abstract

Genome-wide association studies (GWAS) have been highly successful in identifying genomic loci associated with complex traits. However, identification of the causal genes that mediate these associations remains challenging, and many approaches integrating transcriptomic data with GWAS have been proposed. However, there currently exist no computationally scalable methods that integrate total and allele-specific gene expression to maximize power to detect genetic effects on gene expression. Here, we describe a unified framework that is scalable to studies with thousands of samples. Using simulations and data from GTEx, we demonstrate an average power gain equivalent to a 29% increase in sample size for genes with sufficient allele-specific read coverage. We provide a suite of freely available tools, mixQTL, mixFine, and mixPred, that apply this framework for mapping of quantitative trait loci, fine-mapping, and prediction.

## 1 Introduction

Genome-wide association studies (GWAS) have identified tens of thousands of genomic loci associated with complex traits. A large majority of these loci lie in non-coding regions of the genome, which hinders identification of the underlying molecular mechanisms and causal genes. Multiple methods have been developed to integrate GWAS results with expression quantitatite trait loci (eQTLs), to test whether complex trait associations are mediated through regulation of gene expression. Two strategies are commonly employed: 1) association-based approaches including PrediXcan [Gamazon et al., 2015], fusion [Gusev et al., 2016], and smr [Zhu et al., 2016]; and 2) colocalization-based approaches including coloc [Giambartolomei et al., 2014], eCAVIAR [Hormozdiari et al., 2016], and enloc [Wen et al., 2017]. These approaches rely on high-quality eQTL mapping, fine-mapping, and gene expression predictions.

In cis-eQTL analysis, allele-specific expression (ASE), i.e., the relative expression difference between the two haplotypes, captures the genetic effect of nearby variants. ASE provides additional signal to total read count, and several methods have been proposed to combine total and allele-specific read count for QTL mapping, such as TReCASE [Sun, 2012], WASP [Van De Geijn et al., 2015], and RASQUAL [Kumasaka et al., 2016]). However, these methods are computationally too costly to be applied to sample sizes beyond a few hundred and as a result have not been applied to large-scale studies like GTEx, which includes over 17,000 samples across 49 tissues. Recently, two fine-mapping approaches have been proposed utilizing effect size estimates obtained from both ASE and eQTL mapping via meta-analysis [Zou et al., 2019; Wang et al., 2020]. However, no existing methods, to our knowledge, provides a unified framework of total and allele-specific counts with explicit multi-SNP modeling for QTL mapping, fine-mapping, and prediction.

By assuming a log-linear model for transcript expression levels with independent reads from each haplotype and weak genetic effects, as proposed by [Mohammadi et al., 2017], we derive two approximately independent equations for allelic imbalance (read count difference between the two haplotypes) and total read count. This enables us to develop computationally efficient algorithms for cis-QTL mapping, fine-mapping, and prediction. We demonstrate the resulting gain in performance through simulations under a range of different settings, applications to GTEx v8 data [Aguet et al., 2019], and comparisons to a large-scale eQTL meta-analysis from eQTLGen [Võsa et al., 2018].

The software, simulation, data preprocessing, and analysis pipeline can be found at https://github.com/hakyimlab/mixqtl and https://github.com/liangyy/mixqtl-pipeline. A computationally efficient GPU-based implementation of mixQTL has been embedded in tensorQTL https://github.com/broadinstitute/tensorqtl.

## 2 Results

### Overview of the statistical model

To develop a computationally efficient approach that integrates total and allele-specific count data, we assumed multiplicative cis-regulatory effects and noise, similarly to the model proposed in [Mohammadi et al., 2017]. For a given gene, we modeled the haplotypic read count 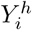, which is the number of reads from haplotype *h* of individual *i* as

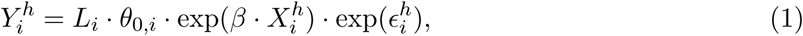

where *L*_*i*_ is the library size for individual *i, θ*_0,*i*_ is the baseline abundance (for a haplotype with the reference allele), exp(*β*) is the cis-regulatory effect (allelic fold change due to the presence of the alternative allele), 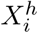 indicates the dosage of the affecting variant (0 if the individual has the reference allele, and 1 if they have the alternative one), and exp 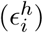 is the multiplicative noise.

Calculating the total read count as the sum of the two haplotypic counts and assuming weak cis-regulatory effects, we derived an approximately linear model for the logorithm of the haplotypic and total read counts (see details in Methods and Supplementary Note 7). In practice, we only observe the allele-specific reads that include a heterozygous site, which is a fraction of the total haplotypic count denoted as 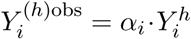. To take this partial readout into account, we modeled the observed total and allele-specific counts as

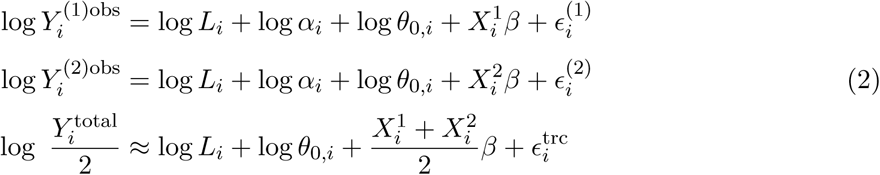

where the error terms are 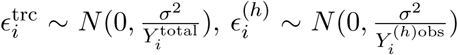 and the errors of the two haplotypes are independent: *ϵ*^(1)^ ╨ *ϵ*^(2)^.

We further simplified the models by combining the two allele-specific counts, defining the baseline abundance variation as a random effect *z*_*i*_ (log *θ*_0,*i*_ = population mean + *z*_*i*_), and dropping 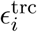 from the total count since this “techinical” noise which scales as the inverse of the read count is small compared to the “biological” variability, *z*_*i*_, (See Methods section and Supplementary Note 10.1) to obtain our final model

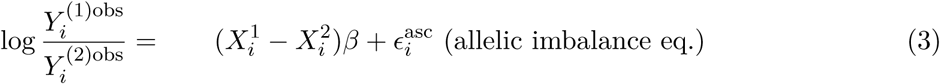

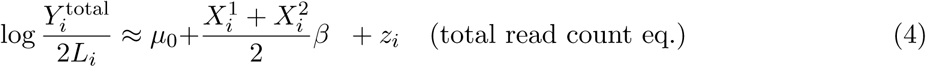

where 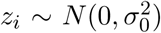 and 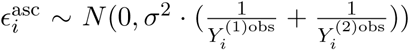 and *z*_*i*_ ╨ *ϵ*^asc^ (baseline abundance is independent of the multiplicative error).

This single SNP model extends to multiple SNPs in a straightforward manner by using a vector of allelic dosages (*X*_1_, …, *X*_*p*_) and genetic effects (*β*_1_, …, *β*_*p*_) instead of the scalar values above. Here, *p* represents the number of genetic variants in the cis-window of the gene under consideration (Supplementary Notes 9 and 11).

For cis-QTL mapping, we took advantage of the approximate independence of the allelic-imbalance and the total read counts in equations (3) and (4), solving them as separate linear regressions (for computational efficiency) and combining the results via inverse-variance weighted meta-analysis. We call this method mixQTL.

For the fine-mapping and prediction problems, we also leveraged the approximate independence of the allelic-imbalance and total read count equations. We used a two-step approach in which we first scale the two equations so that they become independent data points with equal variances. In a second step, we combined these data points into an augmented dataset and applied the existing algorithms SuSiE [Wang et al., 2019] and elastic net [Friedman et al., 2010]. We term these methods mixFine and mixPred, for fine-mapping and prediction, respectively.

### Simulation of total and allele-specific reads

To assess the benefits of this unified framework relative to using total read count or allele-specific expression only, we simulated haplotypic reads according to the framework illustrated in Figure 1, with additional details in Methods (6.7) and Supplementary Notes 12.

**Figure 1:**
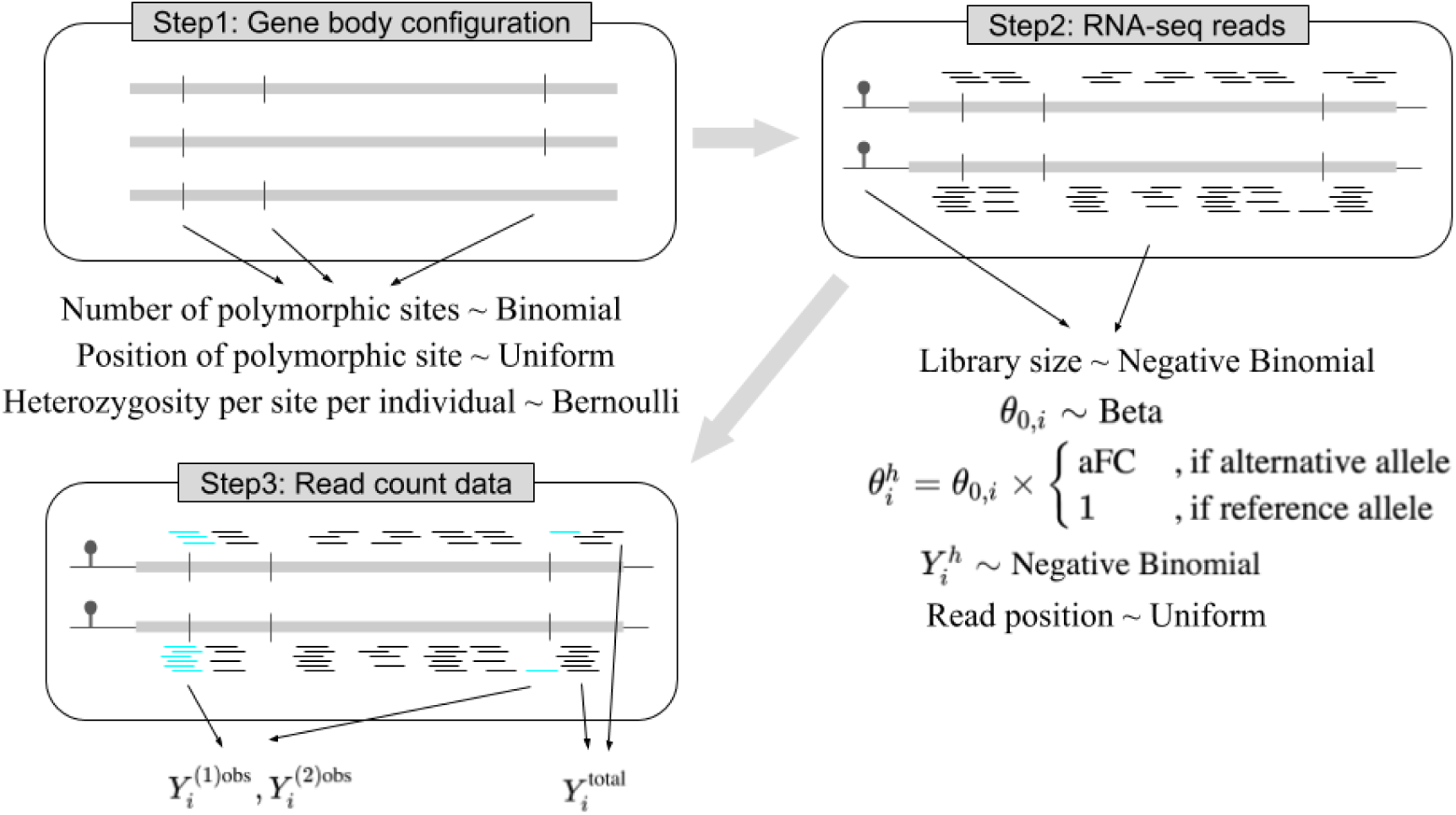
Simulation scheme for total and allele-specific read counts. Step 1 simulates a gene body configuration by first simulating the number of polymorphic sites of the gene followed by positioning these polymorphic sites uniformly across the gene body. For each individual, the actual heterozygosity of these polymorphic sites are drawn from Bernoulli distribution. Step 2 simulates the haplotypic reads by first simulating Negative Binomial library size *L*_*i*_, Beta baseline abundance *θ*_0,*i*_, and the genetic effect *β*. These parameters determine the expected count for each transcript. Then, the actual haplotypic read count 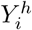 is generated using a Negative Binomial distribution given the expected count where the reads are distributed uniformly across the gene body. In Step 3, the gene-level allele-specific counts 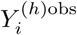 are determined by counting the reads that overlap heterozygous sites, in which aFC is the allelic fold change which equals to *e*^*β*^ in our parameterization. For convenience, we used natural log rather than base 2 log. 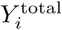 is calculated as the sum of the two haplotypic counts 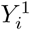 and 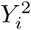.

For all simulation settings, we set an average library size of 94 million reads (to match closely to GTEx v8 library size) and used two expression levels (expected value of *θ*_0,*i*_ in Eq 1): 10 and 1 read per million, corresponding to *θ* = 10^−5^ and 10^−6^. The fraction of allele-specific reads was kept at the similar levels across simulations by using the same distribution of polymorphic sites per individual.

To compare the computational cost of mixQTL to RASQUAL and WASP, we tested their performance on simulated data with 100 samples. As shown in Supplementary Figure S4, type I error and power were similar for all three methods and mixQTL was 10 to 43 times as fast as the others.

### Combining total and allele-specific read counts improves cis-eQTL mapping

To assess the gain in power of combining total and allele-specific read counts, we simulated 200 replicates with allelic fold change varying among 1, 1.01, 1.05, 1.1, 1.25, 1.5, 2, 3. We compared mixQTL with two methods: using either only allele-specific counts (ascQTL) or total counts (trcQTL). See details in Supplementary Note 10.1.

All three methods had calibrated type I errors (Figures 2A and S1). And mixQTL outperformed both trcQTL and ascQTL in all simulation settings, demonstrating the benefits of combining total and allele-specific counts in cis-eQTL mapping (Figures 2B and S2).

**Figure 2:**
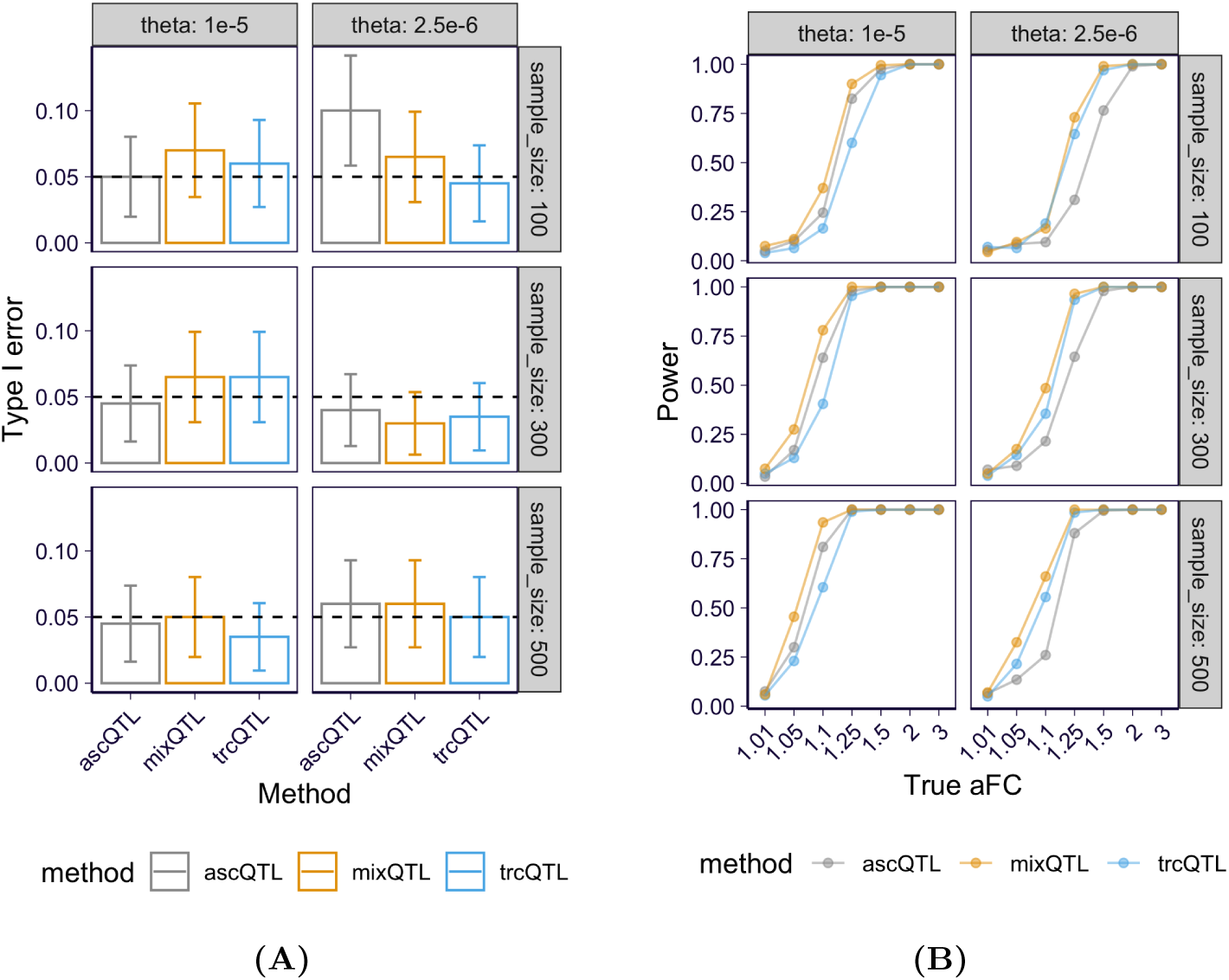
QTL mapping performance for mixQTL and approaches based on either total reads (trcQTL) or allele-specific reads (ascQTL) on simulated data. Each panel presents the results for two relative abundances of the gene, *θ*, and three sample sizes. **(A)** Type I error (y-axis) at a 5% significance level across methods (x-axis). The dashed line represents the desired error rate under the null hypothesis. The error bar indicates the 95% confidence interval (CI) in observed error rate estimated from 200 replicates. **(B)** Power (y-axis) at a 5% significance level across methods under a range of true aFC values (x-axis). Power is defined as the fraction of eQTLs passing the significance threshold.

The degree of improvement varied with the number of reads and sample size. The power of ascQTL was sensitive to the number of allele-specific reads, as expected. As shown in Figure 2B, ascQTL yielded much higher power in the case of relatively large *θ* (on the left) compared with small *θ* (on the right). In contrast, trcQTL was less sensitive to the number of reads observed under the range of read counts in our simulation settings. Such sensitivity differences between ascQTL and trcQTL are consistent with the nature of count data, where the magnitude of the noise is inversely related to the count.

### Combining total and allele-specific read count improves fine-mapping

To mimic LD structure realistically in our simulations, we used the genotypes of European individuals from the 1000 Genomes projects phase 3 [1000 Genomes Project Consortium, 2015] within 1MB cis-windows of 100 randomly selected genes. We applied mixFine and trcFine (which uses total read count only; Supplementary Notes 11.3) to the simulated data and characterized the fine-mapping results with two metrics: 1) power curve, defined as the proportion of detected variants among causal ones versus the number of detected variants, where detection means the variant has posterior inclusion probability (PIP) > some threshold (which is varied to get the desired number of detected SNPs); 2) the size of 95% credible set (CS) which contains the causal variant.

The PIP of both trcFine and mixFine were consistent with the proportion of true causal variants within each bin of 0.1 length (Figure 3A). By combining total and allele-specific reads, mixFine achieved higher power than trcFine (Figures 3B and S5) across all simulation settings. mixFine achieved the highest improvement relative to trcFine at high expression level, *θ*, corresponding to high-quality allele-specific signals. The gain in power decreased with larger sample sizes.

**Figure 3:**
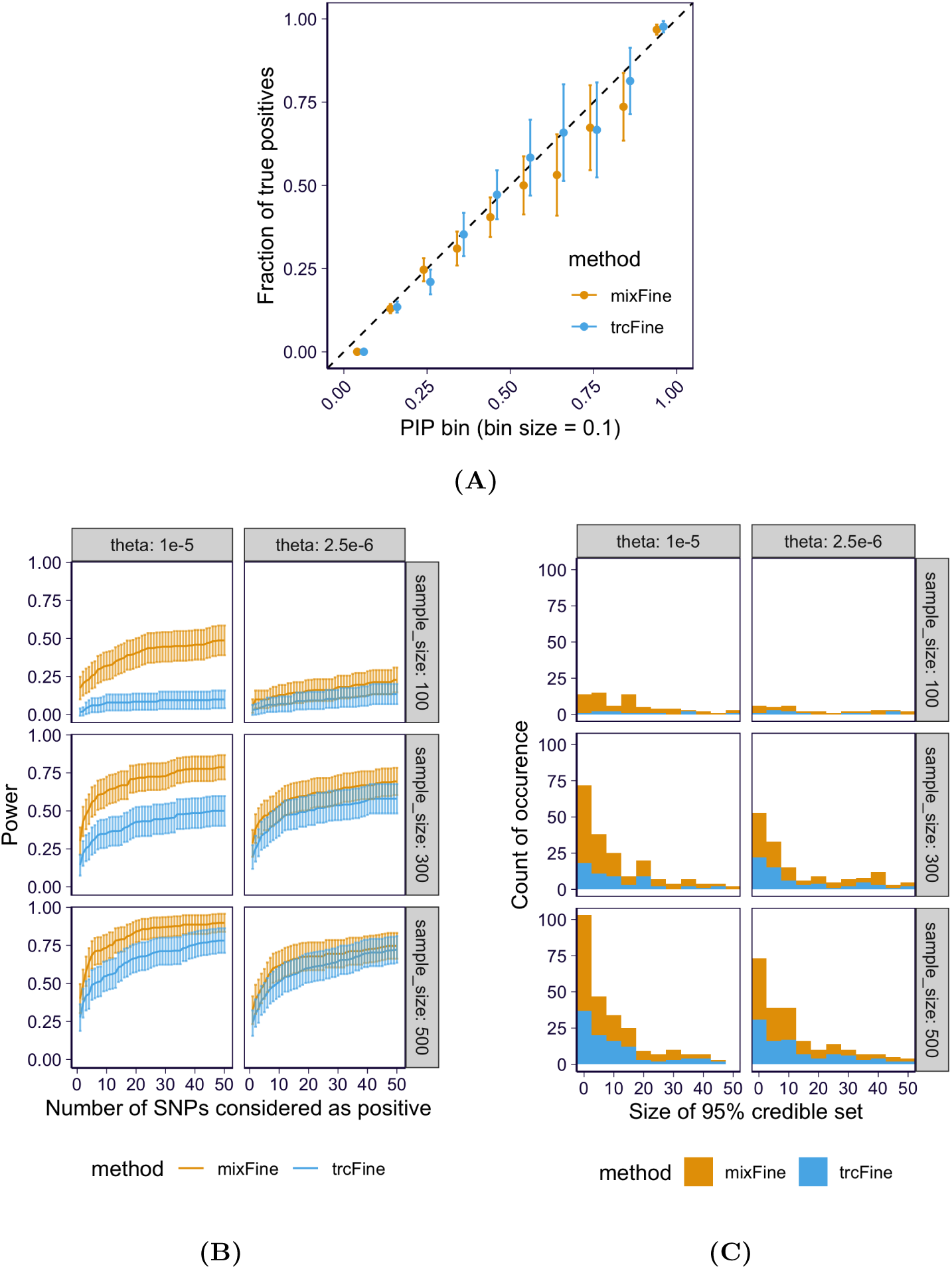
Fine-mapping performance of the combined (mixFine) and total read-based (trcFine) approaches on simulated data. **(A)** The observed true positive rate within SNPs binned by PIP are shown (aggregated across all simulation settings) for both mixFine (orange) and trcFine (blue). **(B)** The power at a PIP cutoff (on y-axis) is plotted against the number of variants passing the PIP cutoff (on x-axis) for mixFine and trcFine. The solid curves indicate the mean power (recall rate) among 100 simulation replicates and the error bars indicate the 95% CI. **(C)** The distribution of the size of 95% CS that contain the causal variant for mixFine and trcFine across all 100 simulation replicates. The counts in each bin are stacked.

The increased power was also reflected in the number and size of 95% CSs containing the true signals. As shown in Figures 3C and S6, mixFine identified more true positive 95% CSs and these 95% CSs were generally smaller than the ones of trcFine demonstrating that mixFine can pinpoint causal SNPs more accurately.

Overall, the combined method was more powerful for identifying causal variants, which is consistent with recent reports [Zou et al., 2019; Wang et al., 2020].

### Combining total and allele-specific read count improves prediction

Using the data from the fine-mapping simulation, we tested the performance of mixPred and trcPred (Supplementary Notes 11.3) on held-out test data. Specifically, we split each simulation replicate into training (4/5) and test (1/5) sets. We trained prediction models using training data and evaluated the prediction performance on test data using Pearson correlation between predicted and true response. For each data set, we repeated the splitting-training-evaluation procedure twice to reduce the stochasticity introduced by splitting.

Overall, mixPred achieved higher prediction accuracy than trcPred (Figure 4 and Supplementary Figure S7 and S8). The gain in performance was more apparent when the expression level *θ* was higher and as a consequence the allele-specific count was larger.

**Figure 4:**
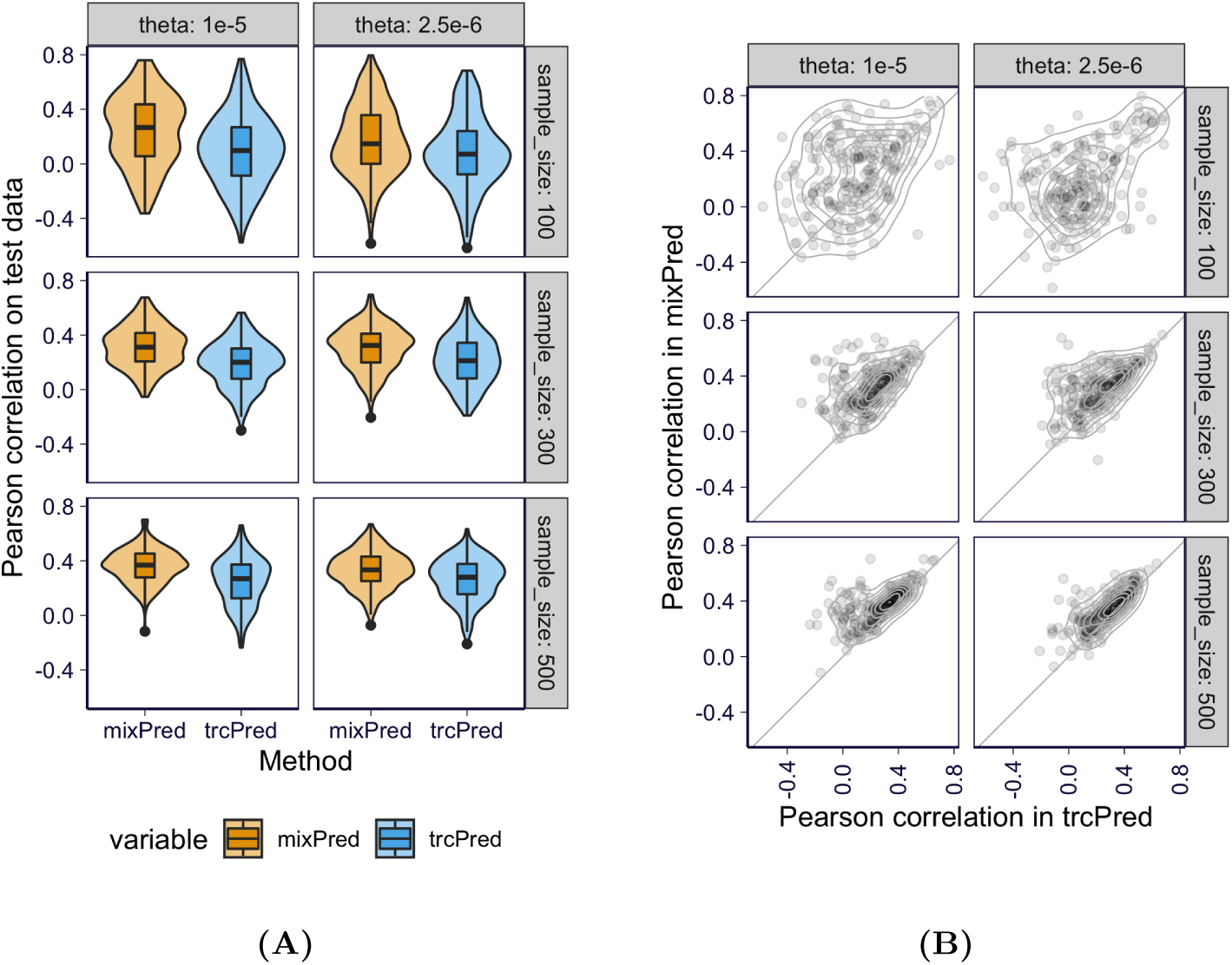
Prediction performance of the combined (mixPred) and total read-based (trcPred) methods on simulated data. **(A)** The overall distribution of Pearson correlations between predicted and observed total count abundance in log-scale, i.e., 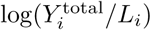, for mixPred (orange) and trcPred (blue) across all data splits are shown. **(B)** For each split, the prediction performance of mixPred (y-axis) is plotted against the prediction performance of trcPred (x-axis).

### mixQTL outperforms standard eQTL mapping in GTEx data

Next, we compared mixQTL to the standard eQTL mapping approach (denoted here simply as eQTL) used by the GTEx consortium [Aguet et al., 2019], using 670 whole blood RNA-seq samples from the v8 release. We included variants within a ±1 Mb cis-window around the transcription start site of each gene, and limited our analysis to genes passing the following two criteria: 1) at least 15 samples having at least 50 allele-specific counts for each haplotype; and 2) at least 500 samples having a total read count of at least 100. 28% of genes passed these filters, corresponding to 5,734 genes in total. For genes with below-threshold allele-specific counts, the calculation is performed using total read counts only, such that all genes considered using the standard approach are also tested in mixQTL. Performance for these genes was similar to the standard eQTL approach (Supplementary Figure S9). We then stratified genes that passed the filtering criteria by their median expression level (read counts) into low, medium, and high expression tertiles.

All three approaches mixQTL, aseQTL, and trcQTL were relatively well-calibrated when permuting data in four randomly selected genes (Supplementary Figure S10). The estimated effect sizes were consistent with allelic fold change estimates from the main GTEx v8 analysis (Supplementary Figure S11).

To further compare the performance of the methods, we used eQTLGen [Võsa et al., 2018], a large-scale meta-analysis of over 30,000 blood samples, as our “ground truth” eQTL discovery reference (Supplementary Notes 14). We selected a random subset of 100,000 variant/gene pairs tested by eQTLGen with FDR < 0.05 as the set of “ground truth” eQTLs. We also selected a random set 100,000 variant/gene pairs with p > 0.50 as a background set of “non-significant” eQTLs. Only 96,660 and 78,691 of the “ground truth” and “non-significant” pairs were found in the GTEx data.

For the “ground truth” eQTLs, mixQTL yielded more significant p-values compared to the standard eQTL, ascQTL, and trcQTL approaches (Fig. 5). The “non-significant” variant/gene pairs showed moderate enrichment for small p-values for all methods (Figure 5B), likely reflecting a combination of false negatives in eQTLGen and potential false positives in our analysis. Overall, we found that mixQTL achieves increased power compared to standard eQTL mapping on real data for the set of genes with sufficient total and allele-specific read counts.

**Figure 5:**
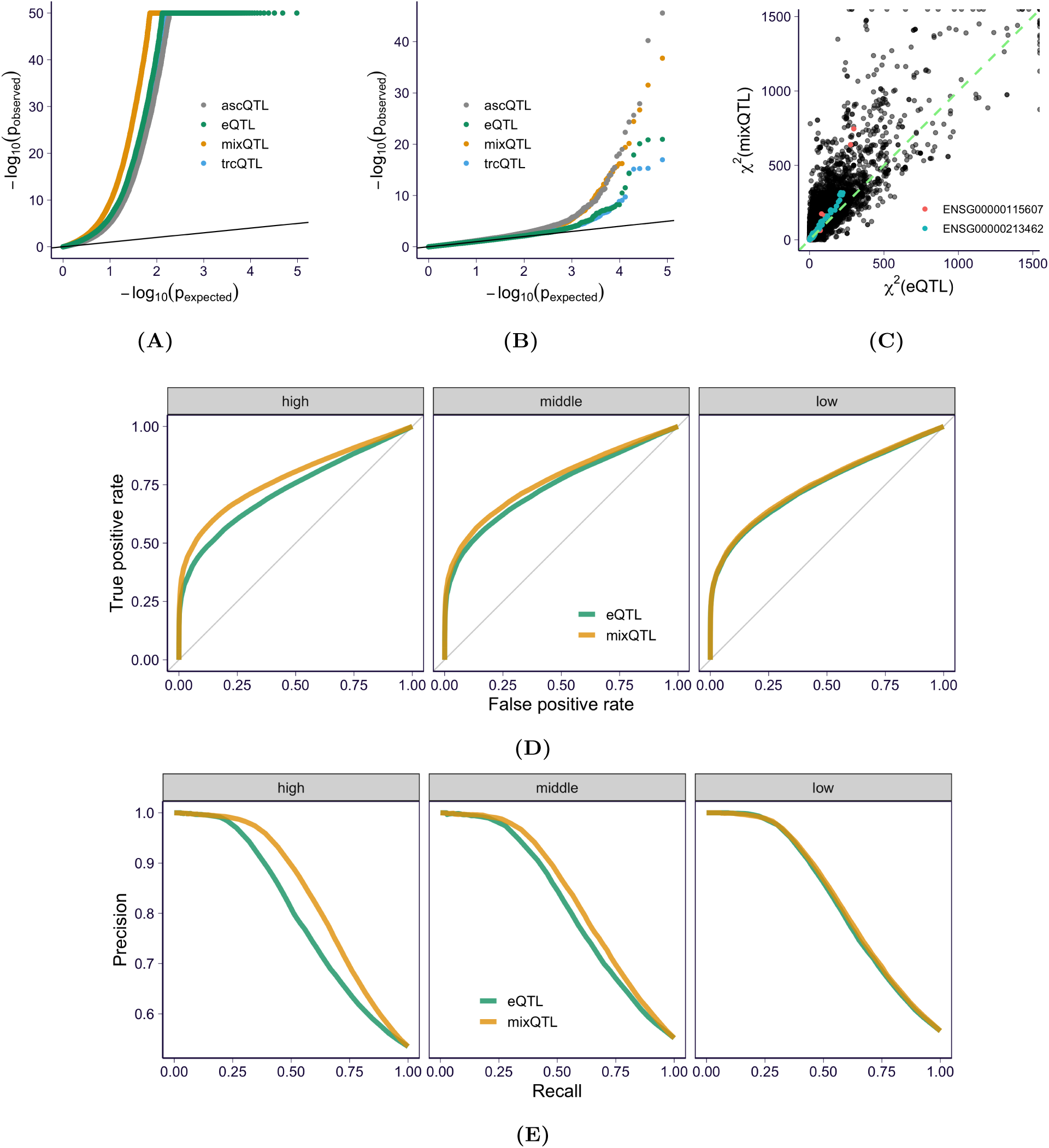
Performance of mixQTL on GTEx v8 whole blood RNA-seq. **(A)** QQ-plot of nominal p-values for a random subset of cis-eQTLs (FDR < 0.05) reported in eQTLGen. **(B)** QQ-plot of nominal p-values for a random subset of variant/gene pairs with p-value > 0.5 in eQTLGen. **(C)** *χ*^2^ statistics from eQTL analysis (x-axis) and mixQTL analysis (y-axis) among a random subset of cis-eQTLs (FDR < 0.05) reported in eQTLGen. The slope indicates the relative effective sample size increase. Two randomly selected genes are highlighted in red and green, respectively. **(D, E)** ROC and PR curves for mixQTL and the standard eQTL method measured in eQTLGen. Each panel shows the results of genes stratified by expression level tertiles.

As an intuitive measure of improved performance, we estimated the effective sample size gain of mixQTL compared to standard eQTL mapping as the median of the ratio between mixQTL *χ*^2^ statistics and eQTL *χ*^2^ statistics. mixQTL showed a 29% increase in effective sample size compared to the standard eQTL mapping approach (Figure 5C).

To account for the trade-off between true and false positive rates, as well as between precision and power, we used receiver operating characteristic (ROC) and precision-recall (PR) curves to compare the performance of mixQTL and standard eQTL approaches using the eQTLGen “ground truth” and “non-significant” eQTLs. We found that mixQTL achieves higher performance in both ROC (Figure 5D) and PR curves (Figure 5E). Consistent with simulation results, this gain is more significant for genes with higher expression levels.

### Fine-mapping and prediction model building in GTEx data

We applied mixFine to the GTEx v8 whole blood RNA-seq data, using the same subset of genes with high expression and allelic counts that were used for QTL mapping above. We compared mixFine to the SuSiE fine-mapping approach [Wang et al., 2019] applied to the inverse normal transformed expression values used for standard eQTL mapping [Aguet et al., 2019]. We corrected for sex, 5 genetic principal components, WGS platform, WGS library prep protocol (PCR), and 60 PEER factors. We refer to the latter as the “standard approach” below for simplicity.

To compare the power of causal variant detection, we performed a subsampling analysis on a random subset of 1,000 genes. First, we defined “consensus SNPs” as the variants with PIP > 0.5 in both mixFine and the “standard approach” using all samples. Similarly, a variant was defined as “top SNP” if it was the most significant variant within the 95% CS for both mixFine and the “standard approach”. Then, we compared how well the “consensus SNPs” and “top SNPs” were detected by mixFine and the standard fine-mapping approach using only a subset of samples. We subsampled to 90%, 80%, …, 30% of samples, and repeated each random subsampling step 10 times.

At each subsampling level, mixFine, on average, detected more “consensus SNPs” than the standard approach (Figure 6A) and performance improved most on the more highly expressed genes (top tertile) (Figure S12). Moreover, mixFine detected “top SNPs” in 95% CSs with average size = 9.6 variants whereas the corresponding 95% CS of standard approach had average size = 13.6 variants (Figure S13). These results indicate that, when sufficient counts are available, mixFine, the multi-SNP model combining total and allele-specific counts, can better pinpoint causal cis-eQTLs than the standard approach on real data.

**Figure 6:**
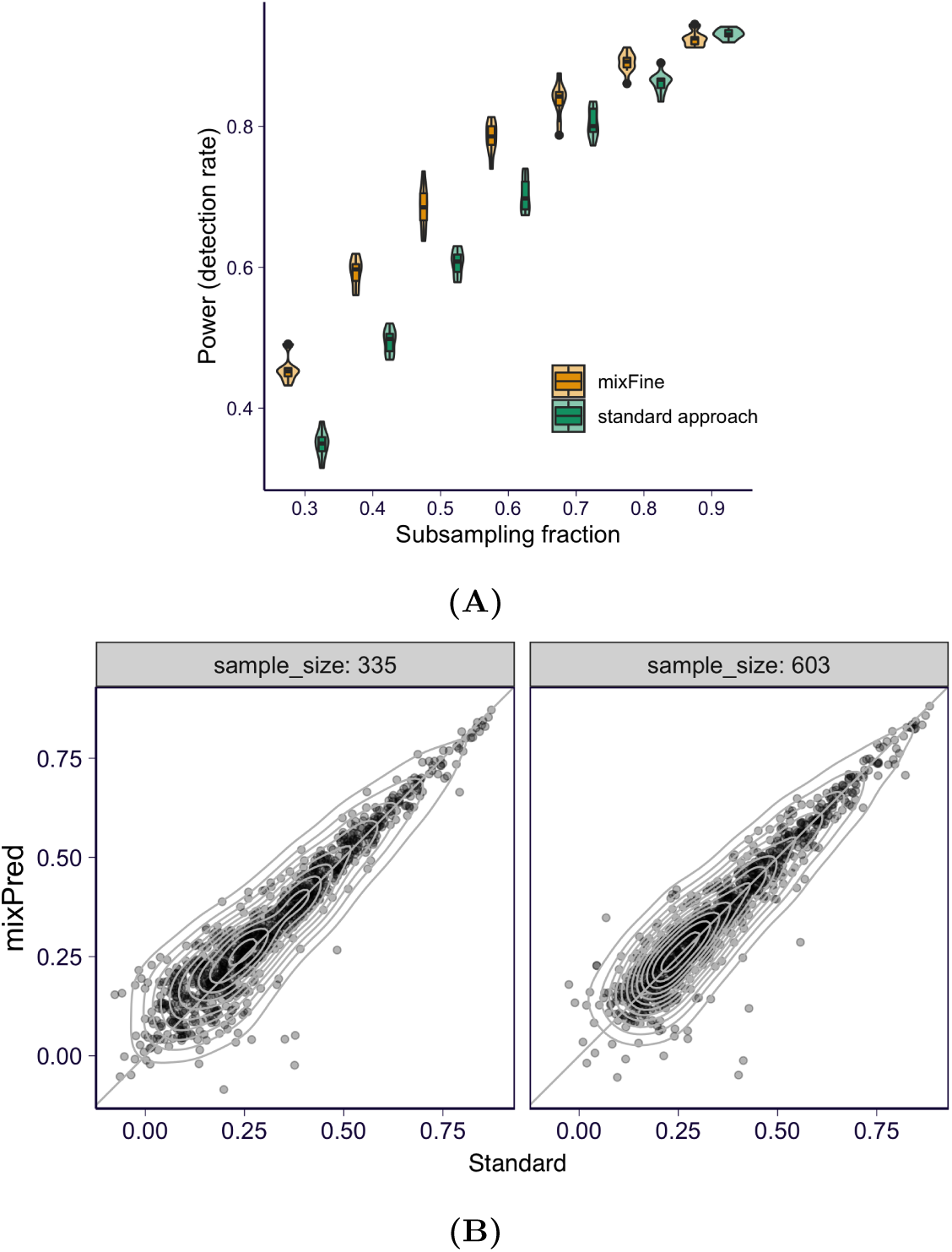
Performance of mixFine and mixPred on GTEx v8 whole blood RNA-seq. **(A)** Fraction of detected “consensus SNPs” as a function of subsampling level, for mixFine and the standard approach. **(B)** Median Pearson correlation across *k* held-out folds for mixPred vs. the standard method, for *k* = 2, corresponds to training sample size = 335 (left panel) and *k* = 10, corresponding to training sample size = 603. Each point corresponds to a gene.

To compare the performance of mixPred and the standard method on real data, we implemented a cross-validated evaluation pipeline where we split the full data into *k* folds. At each fold, we trained the prediction model using the remaining (*k* − 1) folds and evaluated the performance (by Pearson correlation between predicted and observed 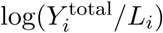) on the held out fold. We applied this evaluation pipeline to mixPred and the standard approach (based on inverse normalized expression) on the same 1000 genes as the subsampling analysis with *k* equals to 2 and 10 (corresponds to sample size = 335 and 603). Both mixPred and the standard approach achieved higher prediction performance as sample size increased, suggesting that sample size was not saturated and was a limiting factor of the prediction performance (Supplementary Figure S14). At the same sample size, we observed, on average, higher performance in mixPred as compared to the standard approach, and the performance gain was more obvious for smaller sample sizes (Figure 6B). These results indicate that mixPred can improve over the standard approach for building prediction models by leveraging allele-specific counts as extra observations.

## 3 Discussion

We proposed a unified framework that integrates both allele-specific and total read counts to estimate genetic cis-regulatory effects, resulting in improved eQTL mapping, fine-mapping, and prediction of gene expression traits. Our suite of tools (mixQTL, mixFine, and mixPred) can be scaled to much larger sample sizes (thousands) due to the underlying log-linear approximation. By assuming multiplicative genetic effects, we transform the observed read counts into two approximately independent quantities: allelic imbalance and total read count. We take advantage of this independence to develop computationally efficient approaches that integrate both allele-specific and total reads.

Specifically, mixQTL estimates the genetic effect separately for the allelic imbalance and the total read counts, and combines the resulting statistics via meta-analysis. These calculations have computationally efficient closed-form solutions, enabling their use in the permutation schemes applied to compute FDR in eQTL mapping [Shabalin, 2012; Ongen et al., 2015; Taylor-Weiner et al., 2019].

Furthermore, the simple multi-SNP extension and the independence of the terms enable use of a two-step inference procedure. In the first step, the allelic imbalance and total read count are scaled so that the error terms have the same variance. And in the second step, given their approximate independence, the pair of equations (from allelic imbalance and total counts) can simply be input into existing fine-mapping and prediction algorithms. We showed through extensive simulations and applications to GTEx v8 data that our suite of methods outperforms current methods that use only total read count. Given the straightforward extension of current approaches with the models proposed here, as well as their computational efficiency, we anticipate that combining total and allele-specific read counts will find widespread use for eQTL mapping, fine-mapping, and prediction of gene expression.

## 4 Acknowledgement

We used data from the GTEx project (dbGap accession number phs000424.v8.p2). The Genotype-Tissue Expression (GTEx) Project was supported by the Common Fund of the Office of the Director of the National Institutes of Health (commonfund.nih.gov/GTEx). Additional funds were provided by the NCI, NHGRI, NHLBI, NIDA, NIMH, and NINDS. Donors were enrolled at Biospecimen Source Sites funded by NCI Leidos Biomedical Research, Inc. subcontracts to the National Disease Research Interchange (10XS170), Roswell Park Cancer Institute (10XS171), and Science Care, Inc. (X10S172). The Laboratory, Data Analysis, and Coordinating Center (LDACC) was funded through a contract (HHSN268201000029C) to the The Broad Institute, Inc. Biorepository operations were funded through a Leidos Biomedical Research, Inc. sub-contract to Van Andel Research Institute (10ST1035). Additional data repository and project management were provided by Leidos Biomedical Research, Inc.(HHSN261200800001E). The Brain Bank was supported supplements to University of Miami grant DA006227. Statistical Methods development grants were made to the University of Geneva (MH090941 & MH101814), the University of Chicago (MH090951,MH090937, MH101825, & MH101820), the University of North Carolina - Chapel Hill (MH090936), North Carolina State University (MH101819),Harvard University (MH090948), Stanford University (MH101782), Washington University (MH101810), and to the University of Pennsylvania (MH101822). The datasets used for the analyses described in this manuscript were obtained from dbGaP at http://www.ncbi.nlm.nih.gov/gap through dbGaP accession number phs000NNN.vN.pN

HKI, YL, and AB were partially funded by R01MH10766 and P30 DK20595 (Diabetes Research and Training Center).

## Code and data availability

Software mixQTL, mixFine, and mixPred https://github.com/hakyimlab/mixqtl

Reproducible pipeline https://github.com/liangyy/mixqtl-pipeline

GPU-based implementation embedded in tensorQTL https://github.com/broadinstitute/tensorqtl.

## 6 Material and Methods

### 6.1 Notation and terminology

**Table 1:** Summary of notation and terminology used in the paper. The **Description** column contains a brief definition of each **Notation**, and the **Synonym in text** column contains the corresponding terminology used in the text. The **Observable** column indicates whether the entity is an observable variable or not. (†, ⋆: expression level does not strictly refer to 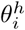 or E(*θ*_0,*i*_), but more generally to the abundance of the gene transcripts relative to the transcriptome.)

| Notation | Description | Synonym in text | Observable |
| --- | --- | --- | --- |
| $i$ | Individual index. | - | - |
| $h$ | Haplotype index, with $h = 1, 2$ for diploid. | - | - |
| $X_i^h$ | Alternative allele count (0 or 1) of the variant linking to the gene haplotype $h$. | allelic dosage | Yes |
| $L_i$ | The total number of reads in the RNA-seq library. | library size | Yes |
| $Y_i^h$ | Count of reads originated from gene haplotype $h$. | haplotypic (read) count | No |
| $Y_i^{(h)\text{obs}}$ | Allele-specific read count that gets aligned to the gene haplotype $h$. | allele-specific (read) count | Yes |
| $Y_i^{\text{total}}$ | Total count of reads originated from any of the two gene haplotypes (sum). | total (read) count | Yes |
| $\theta_{0,i}$ | The abundance of the gene haplotype relative to the total transcriptome when the linked causal variants are all in reference alleles | baseline (relative) abundance | No |
| $\theta_i^h$ | The abundance of the gene haplotype $h$ relative to the total transcriptome in individual $i$ | (relative) abundance; expression level<sup>†</sup> | No |
| $\beta$ | The log fold change of gene haplotype abundance when linking to alternative allele relative the reference allele | allelic fold change (aFC) in natural log scale | No |
| $\frac{Y_i^{(1)\text{obs}}}{Y_i^{(2)\text{obs}}}$ | The ratio of the allele-specific counts between two haplotypes | allelic imbalance | Yes |
| $Y_i^{\text{trc}}$ | Shorthand of the term $\log \frac{Y_i^{\text{total}}}{2L_i}$. | - | - |
| $Y_i^{\text{asc}}$ | Shorthand of the term $\log \frac{Y_i^{(1)\text{obs}}}{Y_i^{(2)\text{obs}}}$ | - | - |
| $\theta$ | Only used in simulation where $\theta = E(\theta_{0,i})$ | expression level* | - |

### 6.2 Statistical model of cis-regulation

For individual *i*, let 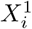 and 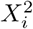 be the number of alternative alleles in each of the two haplotypes at the variant of interest. Let 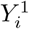 and 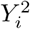 be the number of reads mapped to each of the two haplotypes (i.e., haplotypic counts; in practice, these quantities are unobserved) and *L*_*i*_ the library size for individual *i*. As proposed in [Mohammadi et al., 2017], we use the concept of allelic fold change (aFC) to represent the genetic effect on cis-expression. We denote *θ*_0,*i*_ as the baseline abundance of the transcripts originating from each of the gene haplotype without considering genetic effect. Let *β* be the genetic effect of a variant of interest, which is defined as the log fold change relative to the reference allele. Then, the transcript abundance of each haplotype *h* after accounting for the genetic effect is 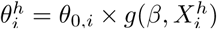 where 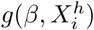 is *e*^*β*^ if 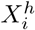 is the alternative allele; otherwise 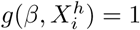. We model read count 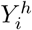 as

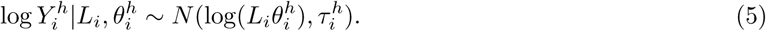

In an RNA-seq experiment, a fraction of reads contribute to allele-specific read counts. Let *α*_*i*_ denote the fraction of allele-specific reads in individual *i*, which depends on the number of heterozygous sites within the transcript. Instead of observing haplotypic counts 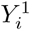 and 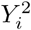, we observe total read count 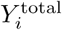 and gene-level allele-specific read counts 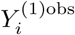 and 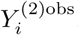. Similarly, we further assume that the baseline abundance of allele-specific reads per haplotype is *θ*_0,*i*_ × *α*_*i*_, so we have

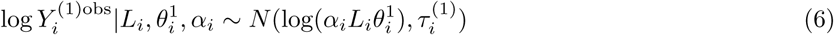

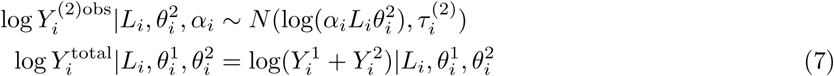

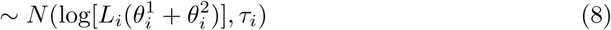

### 6.3 Linearizing the model by approximation

Based on the model described in Section 6.2 along with approximations under weak effect assumptions, we propose the following linear mixed effects model (see Supplementary Notes 8 for derivation):

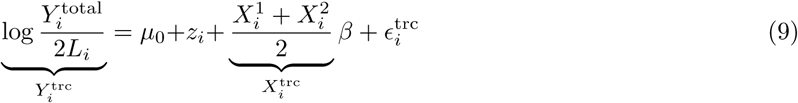

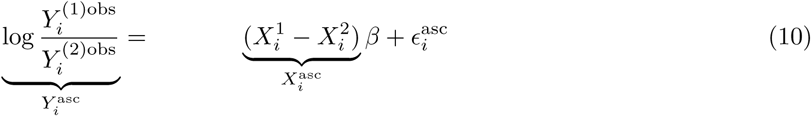

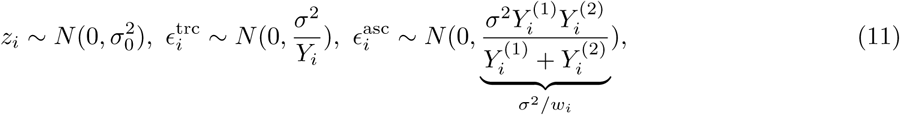

where *z*_*i*_ is the individual-level random effect capturing the between-individual variation of *θ*_*i*,0_. Notice that the individual-level random effect cancels out when we take the difference between the two log-scale allele-specific read counts (i.e., allelic imbalance in log-scale). The scaling of *ϵ*^trc^ and *ϵ*^asc^ in Eq 11 is to ensure that variance of read count scales linearly with the magnitude of read count (see Supplementary Notes 7.2). In other words, this model ensures Var(*Y*) ≈ constant × E(*Y*), such that over-dispersion is implicitly taken into account.

Since *σ*^2^*/Y*_*i*_ is typically much smaller than 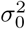 (see Supplementary Notes 10), we can further simplify Eq 9, 10 as

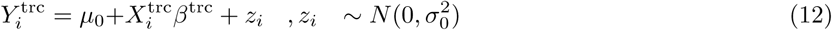

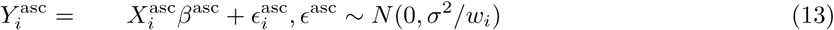

Eqs 12, 13 are applicable to both single-SNP and multi-SNP scenarios. In the single-SNP case, *X*_*i*_ and *β* are scalars, and in the multi-SNP case, *X*_*i*_ and *β* are vectors including all SNPs within the cis-window (see Supplementary Notes 9).

### 6.4 Numerically efficient QTL mapping leveraging approximate independence of allelic imbalance and total read count

The likelihood function corresponding to the proposed model in Eqs 12, 13 takes the form

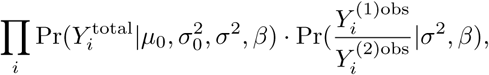

factoring into total read count and allelic imbalance components. (see Supplementary Notes 8.2). This means that the likelihood for total read count and the ratio of allele-specific read counts provide approximately independent information on *β*, and enables us to solve each component separately and combine the results via meta-analysis (standard approach with independent studies [Evangelou and Ioannidis, 2013]). Specifically, we fit *β*^trc^ and *β*^asc^ using total and allele-specific observations as two separate linear regression problems, and meta-analyze the results using inverse-variance weighting (see details in Supplementary Notes 10.2).

### 6.5 Two-step inference procedure for multi-SNP model

The prediction and fine-mapping problems both rely on the linearized model Eq 12, 13, but with different objectives. For prediction, the objective is to find the best predictor, whereas for fine-mapping, the objectiveis to infer whether *β*_*k*_ is non-zero. Existing solvers for both prediction and fine-mapping use total read information only and assume that data (*X, y*) follow the model *y* = *Xβ* + *ϵ*, where the noise term *ϵ* is independent across the rows of the data matrix. We will refer to this model as the ‘canonical’ linear model. We propose a two-step inference procedure that first processes the data such that it approximates *y* = *Xβ* + *ϵ*, and then uses existing solvers for prediction and fine-mapping problems, respectively.

For the first step, we process total and allele-specific reads separately to fit the ‘canonical’ linear model. Specifically, we estimate *σ*^2^ from (*Y* ^asc^, *X*^asc^) based on Eq 13 using elastic net with cross-validation. And similarly, based on Eq 12, we estimate 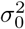 from (*Y* ^trc^, *X*^trc^) and obtain the intercept *µ*_0_ by running fine-mapping with (*Y* ^trc^, *X*^trc^). Then, we shift *Y*^trc^ by 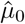 and scale (*Y* ^trc^, *X*^trc^) by 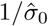. And similarly, we scale (*Y* ^asc^, *X*^asc^) by 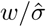. These linear transformations ensure that the transformed 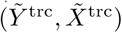 and 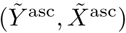 both approximately follow *Y* = *Xβ* + *ϵ*. The implementation details are described in Supplementary Notes 11. At the second step, we concatenate the transformed data from both total and allele-specific read counts as 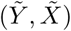, which is compatible with existing solvers for prediction and fine-mapping problems.

### 6.6 Adjusting for covariates

When analyzing real data, we need to take covariates such as sex, batch effect, population stratification into account. Here, we adapt the procedure which has been proposed previously [Mohammadi et al., 2017]. We regress out the effect of covariates beforehand and use the residual as the response in both QTL mapping and fitting multi-SNP model. Specifically, let *c*_1_, *…, c*_*K*_ denote the *K* covariates to be considered. We first regress *Y* ^trc^ against *c*_1_, *…, c*_*K*_ jointly and select the covariates with nominally significant coefficients (p < 0.05). Then we regress *Y* ^trc^ against the selected covariates jointly and set the residuals as the adjusted *Y* ^trc^ for QTL mapping and multi-SNP inference downstream.

### 6.7 Simulation scheme

We simulate RNA-seq reads with total and allele-specific readouts as sketched in three steps in Figure 1. In step 1, we specify, for each individual *i*, the position of heterozygous sites within the gene body. The expected read count from each haplotype transcripts, 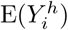, is determined by the RNA-seq library size *L*_*i*_, the baseline abundance of the transcript *θ*_0,*i*_, and the genetic effect *β*. In step 2, given the expected haplotypic count, we draw 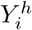 from Negative Binomial to model the variation among count data. In step 3, we position the reads randomly along the gene body and readout observed allele-specific count 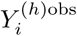 by counting the number of reads overlapping heterozygous sites simulated in step 1. The total read count readout is 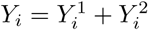, which is independent of the number of heterozygous sites.

To survey a wide range of parameters, we simulate data with a grid of parameters. We vary sample size among 100, 200, …, 500. At library size around 90 million, we vary the level of *θ*_0,*i*_ to cover the gene with different expression levels, among 5 × 10^−5^, 2.5 × 10^−5^, 1 × 10^−5^, 2.5 × 10^−6^, 1 × 10^−6^. The genetic effect, aFC, is set to 1 (null), 1.01, 1.05, 1.1, 1.25, 1.5, 2, 3 in the single SNP model. For the multi-SNP scenario, we set the number of causal SNPs between 1 and 3 with heritability from 0.2 to 0.55. The number of polymorphic sites within the gene body is centered around 10 with minor allele frequency from 0.05 to 0.3. A detailed description and parameter settings are provided in the Supplementary Notes 12.

### 6.8 Analysis of GTEx v8 data

We downloaded the phased genotypes, total read count matrix, and variant-level allele-specific read counts in whole blood from GTEx release 8 [Aguet et al., 2019] via dbGaP (accession number phs000424.v8.p1). To obtain gene-level read counts, we summed over allele-specific counts at all the heterozygous sites for each gene haplotype. We also obtained library size, sex, and genotype PCs from GTEx v8. For comparisons with the inverse normalization-based approach, we also downloaded normalized expression matrices.

Similarly to the GTEx v8 report [Aguet et al., 2019], we restricted the analysis to the cis-regulatory window defined as 1Mbp up/downstream of the transcription start site of each gene.

To obtain the PEER factors for mixQTL analysis, we ran `peertool` [Stegle et al., 2010] on matrix with entry 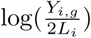 for individual *i* and gene *g* (impute value by k-nearest neighbor if *Y*_*i,g*_ is zero which is done by `impute::impute.knn` in R).

We considered very large allele-specific counts to be outliers likely due to alignment artifacts and removed individuals with allele-specific read counts greater than 1000. To further limit the undue influence of large count outliers on the estimated log fold-change, 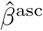, we set the largest weight 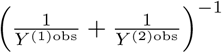to be at most *K* fold to the smallest one, where *K* = min(10, sample size*/*10).

Specific analysis focused on high or low expression were performed with different gene filtering criteria as stated in the Results section.

## Supplementary Figures

**Supplementary Fig. S1.**
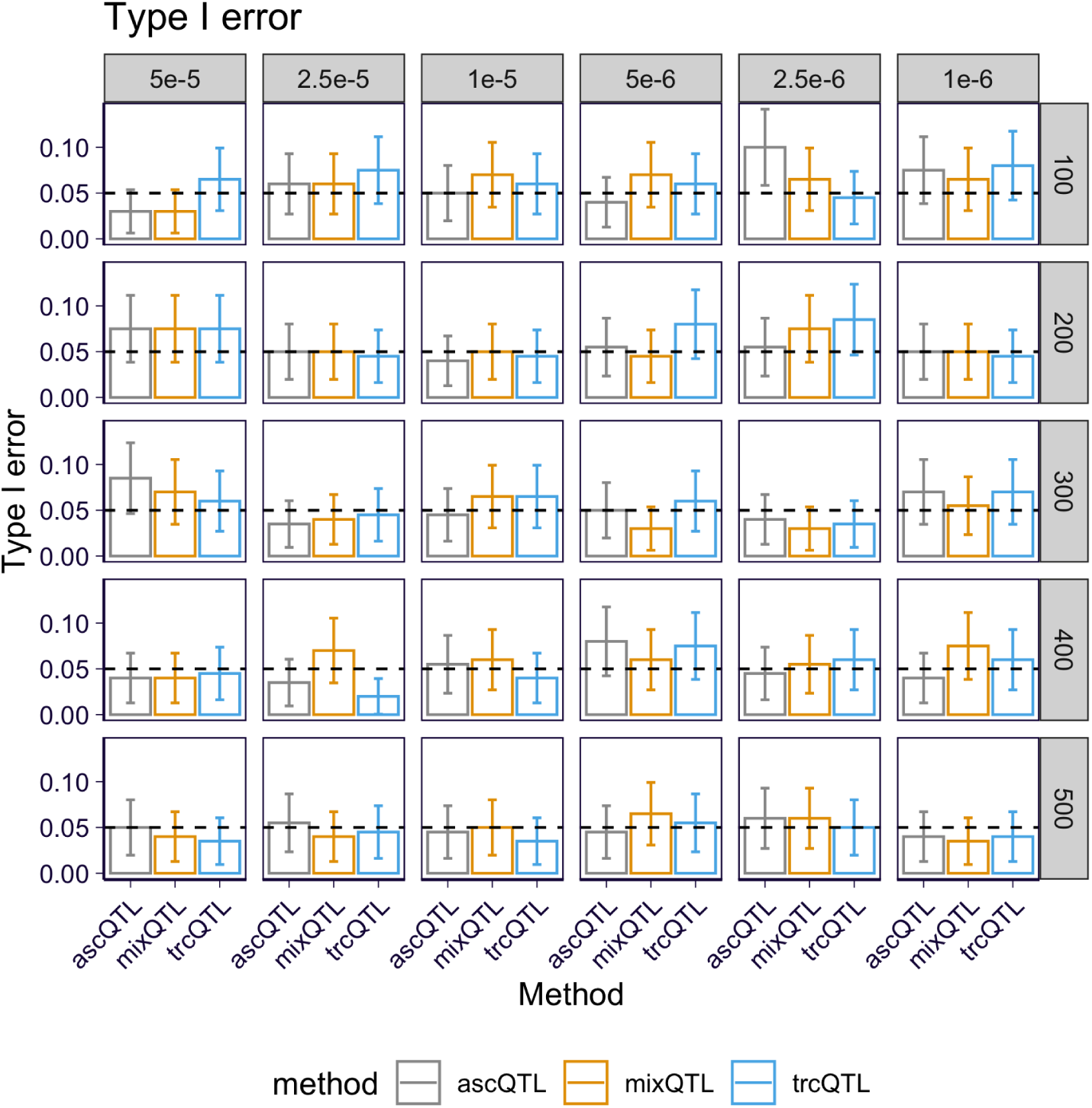
Type I error of mixQTL, ascQTL, and trcQTL on the full grid of simulations. Each panel shows results on data simulated under a pair of *θ* (by column) and sample size (by row).

**Supplementary Fig. S2.**
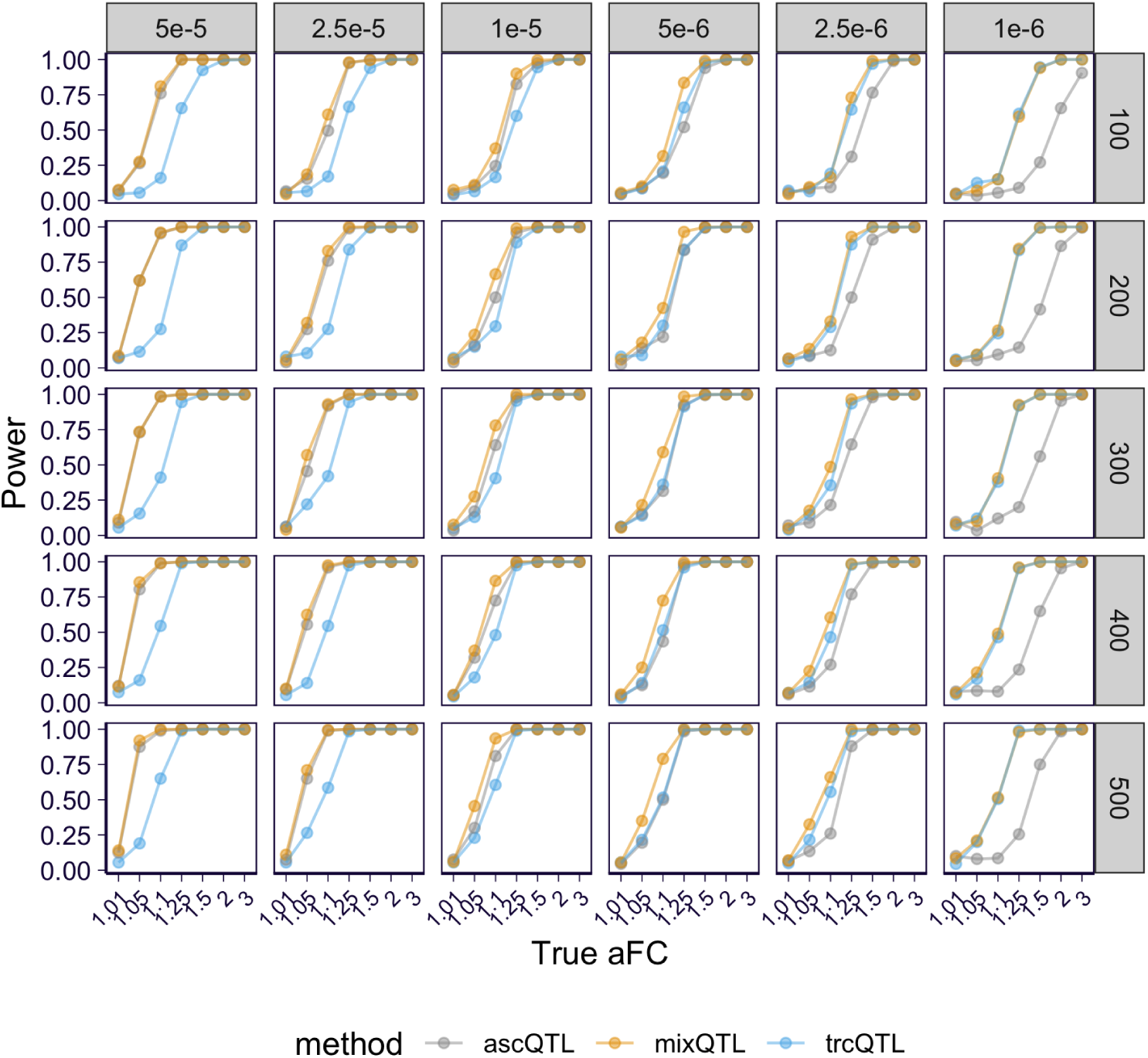
Power of mixQTL, ascQTL, and trcQTL on the full grid of simulations. Each panel shows results on data simulated under a pair of *θ* (by column) and sample size (by row).

**Supplementary Fig. S3.**
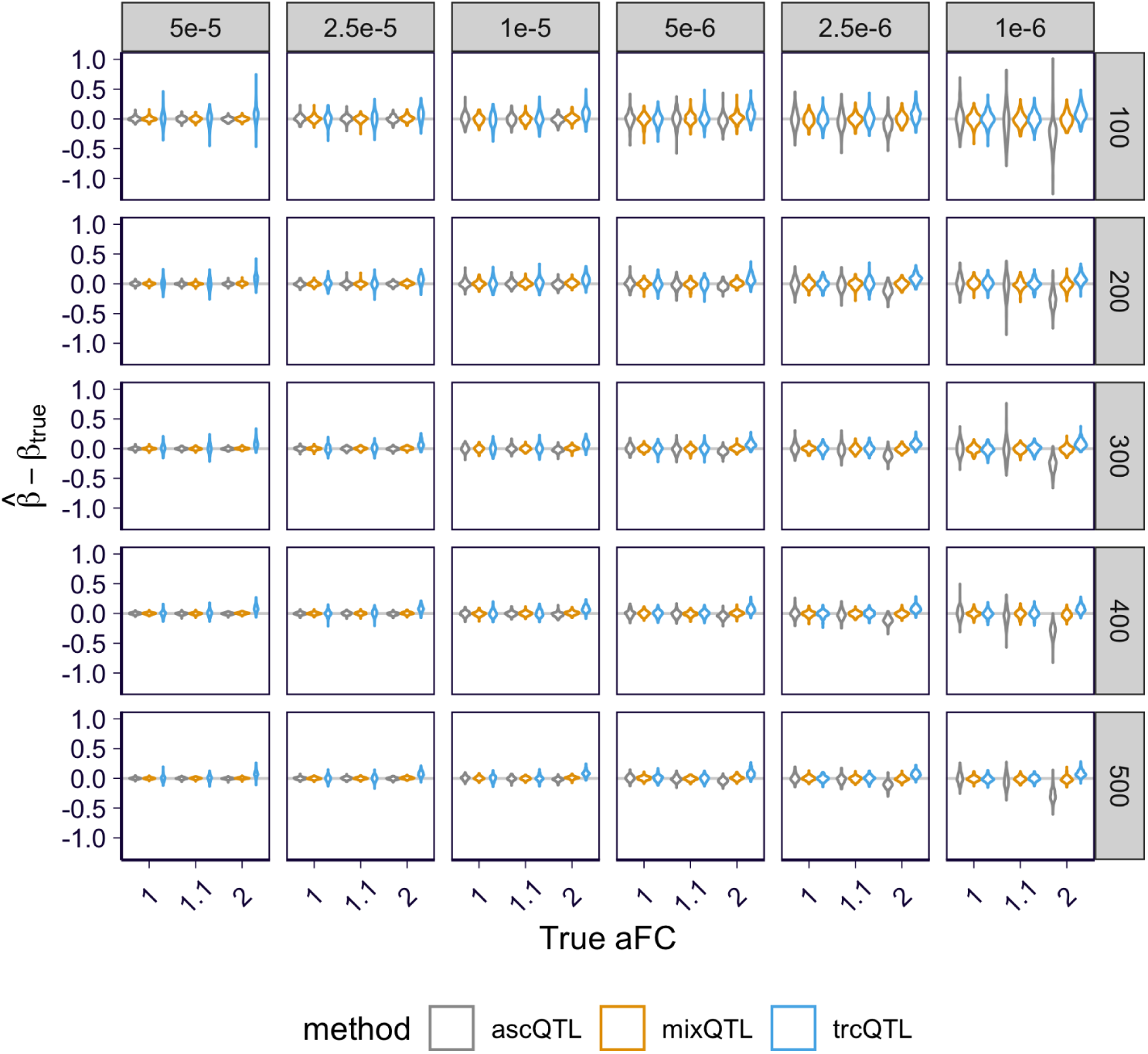
Difference between 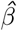 and true *β* of mixQTL, ascQTL, and trcQTL on the full grid of simulations. Each panel shows results on data simulated under a pair of *θ* (by column) and sample size (by row).

**Supplementary Fig. S4.**
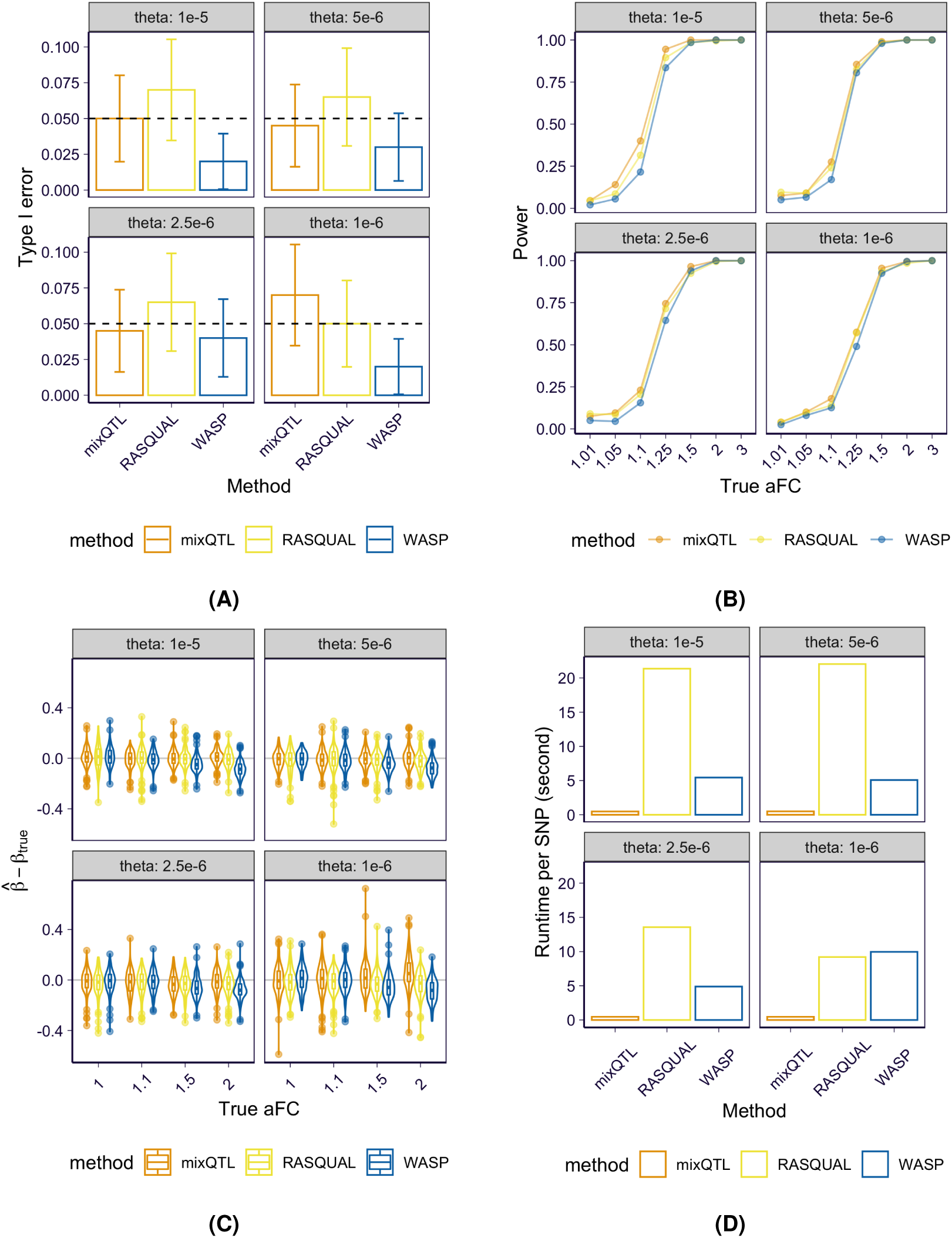
Performance of WASP, RASQUAL, and mixQTL on simulated data. Here, we show results (by panel) for a range of read depths: *θ* = 1 × 10^−5^, 5 × 10^−6^, 2.5 × 10^−6^, 1 × 10^−6^ (defined as the average baseline abundance in the simulation that controls the number of reads) with sample size equal to 100. **(A)** Type I errors computed from 200 replicates. **(B)** Power (y-axis) calculated at *α* = 0.05 for a range of true aFC values (x-axis). **(C)** Difference between estimated aFC and true aFC for a range of true effect sizes (x-axis). **(D)** Per-test computation times. The computation of overdispersion parameters was included.

**Supplementary Fig. S5.**
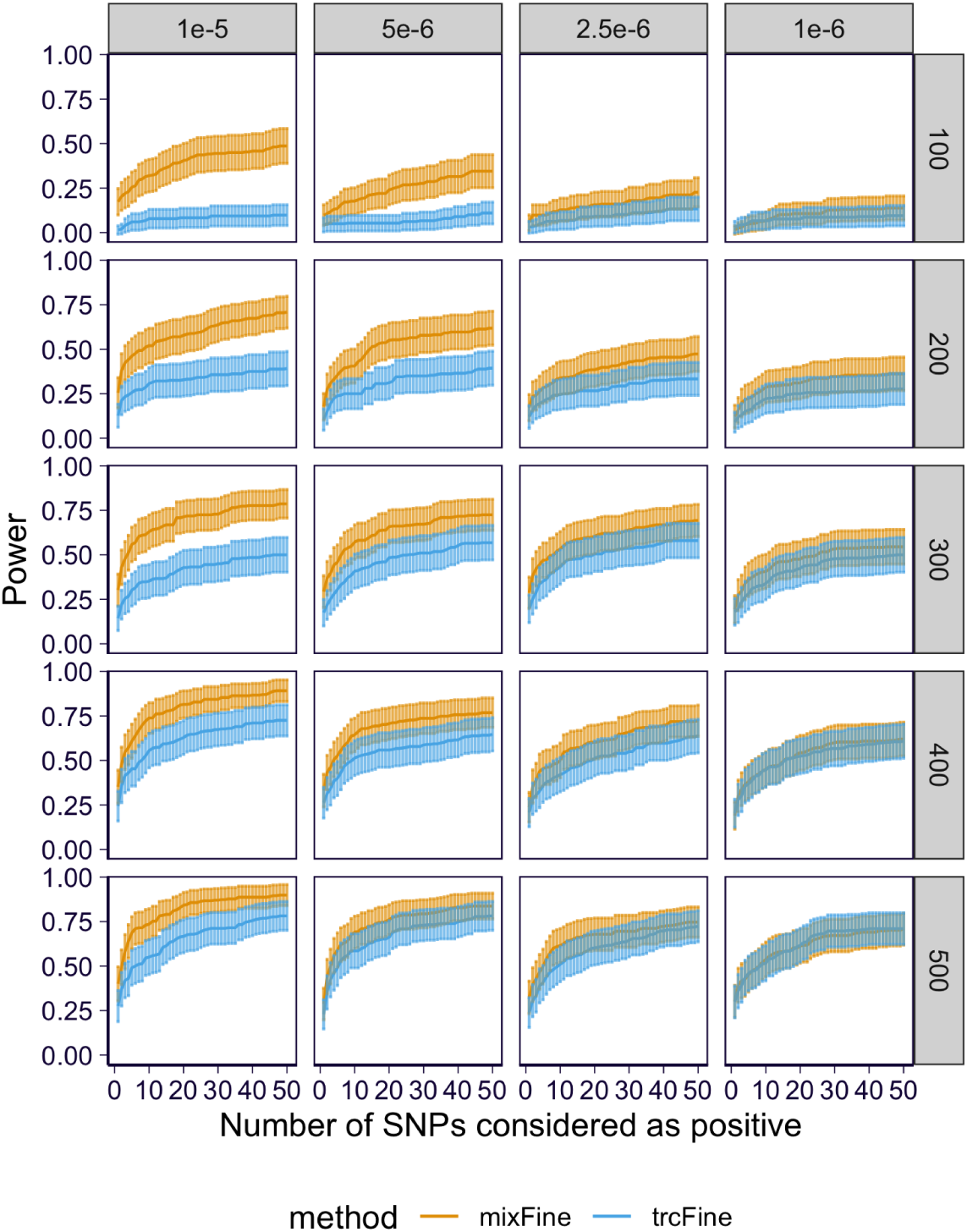
Power curves of mixFine and trcFine on the full grid of simulations. Each panel shows results on data simulated under a pair of *θ* (by column) and sample size (by row).

**Supplementary Fig. S6.**
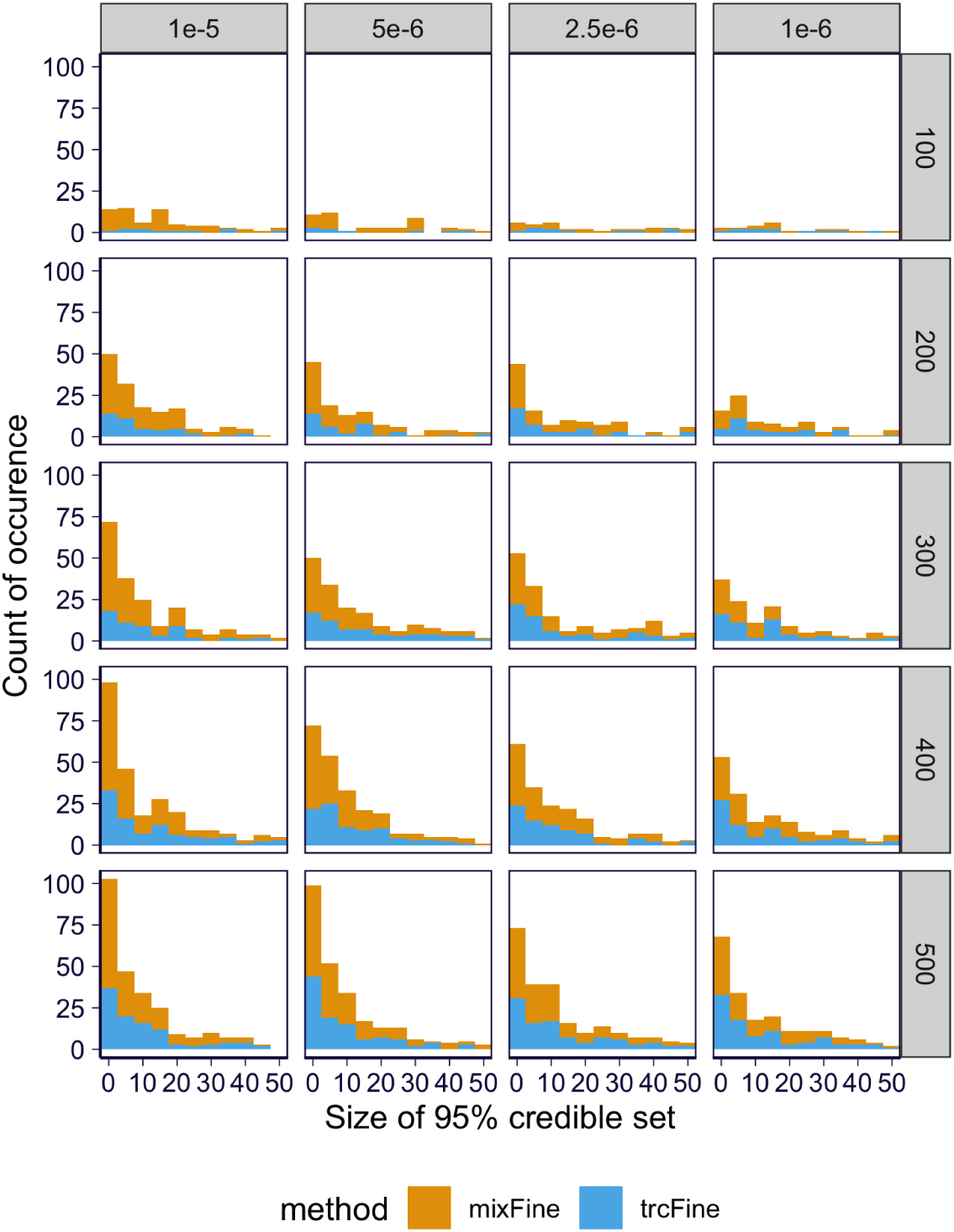
Distribution of the positive 95% CS’s which contain causal variants in mixFine and trcFine on the full grid of simulations. Each panel shows results on data simulated under a pair of *θ* (by column) and sample size (by row).

**Supplementary Fig. S7.**
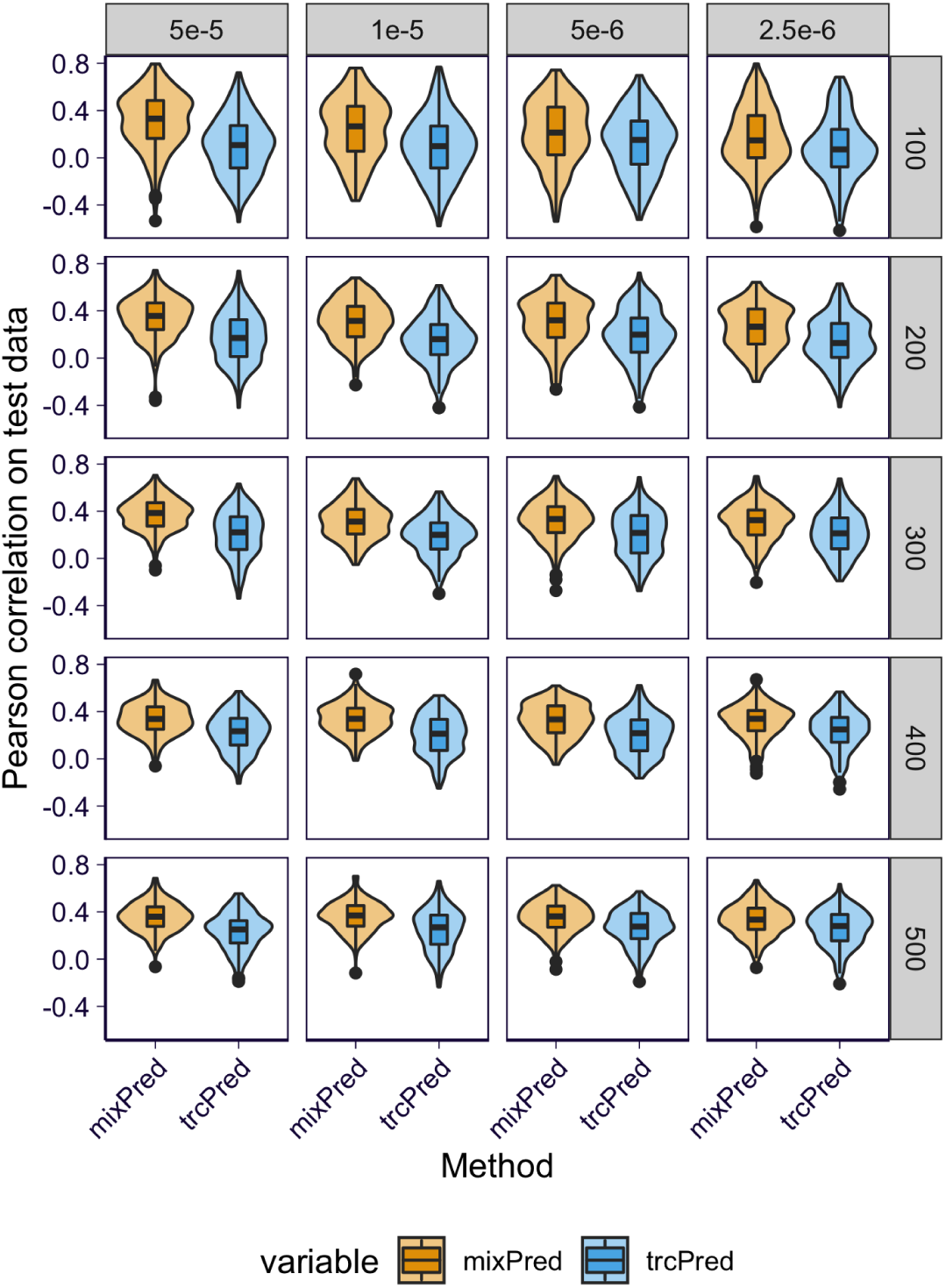
Distribution of Pearson correlations between predicted and observed expression level (in the scale 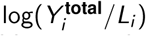) for mixPred and trcPred on the full grid of simulations. Correlation is calculated on held-out test data. Each panel shows results on data simulated under a pair of *θ* (by column) and sample size (by row).

**Supplementary Fig. S8.**
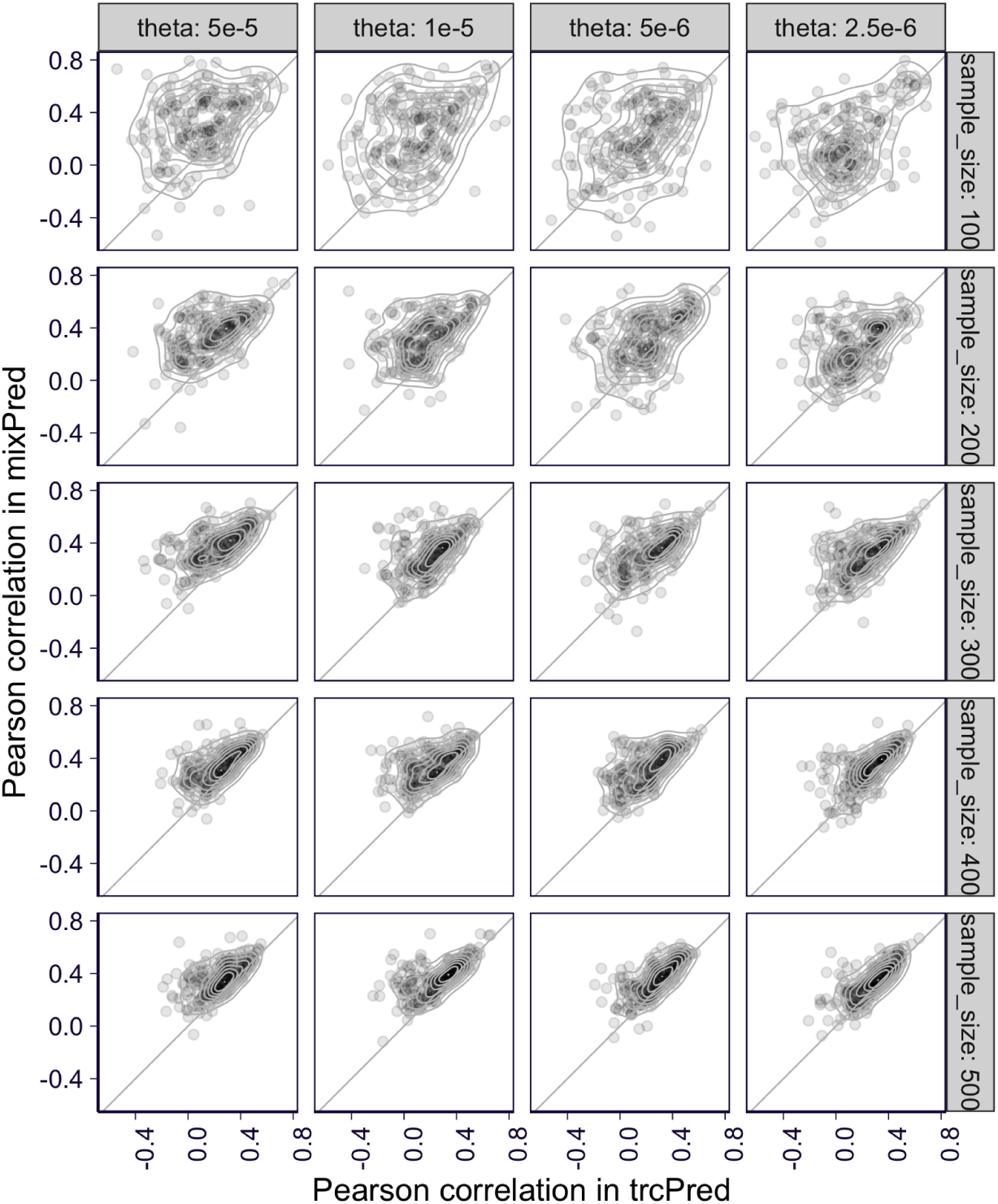
Pairwise comparison of prediction performance of mixPred and trcPred on the full grid of simulations. Correlation of predicted versus observed expression level (in the scale 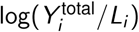) is calculated on held-out test data. The prediction performance of mixPred (y-axis) is plotted against the prediction performance of trcPred (x-axis) for each split. Each panel shows results on data simulated under a pair of *θ* (by column) and sample size (by row).

**Supplementary Fig. S9.**
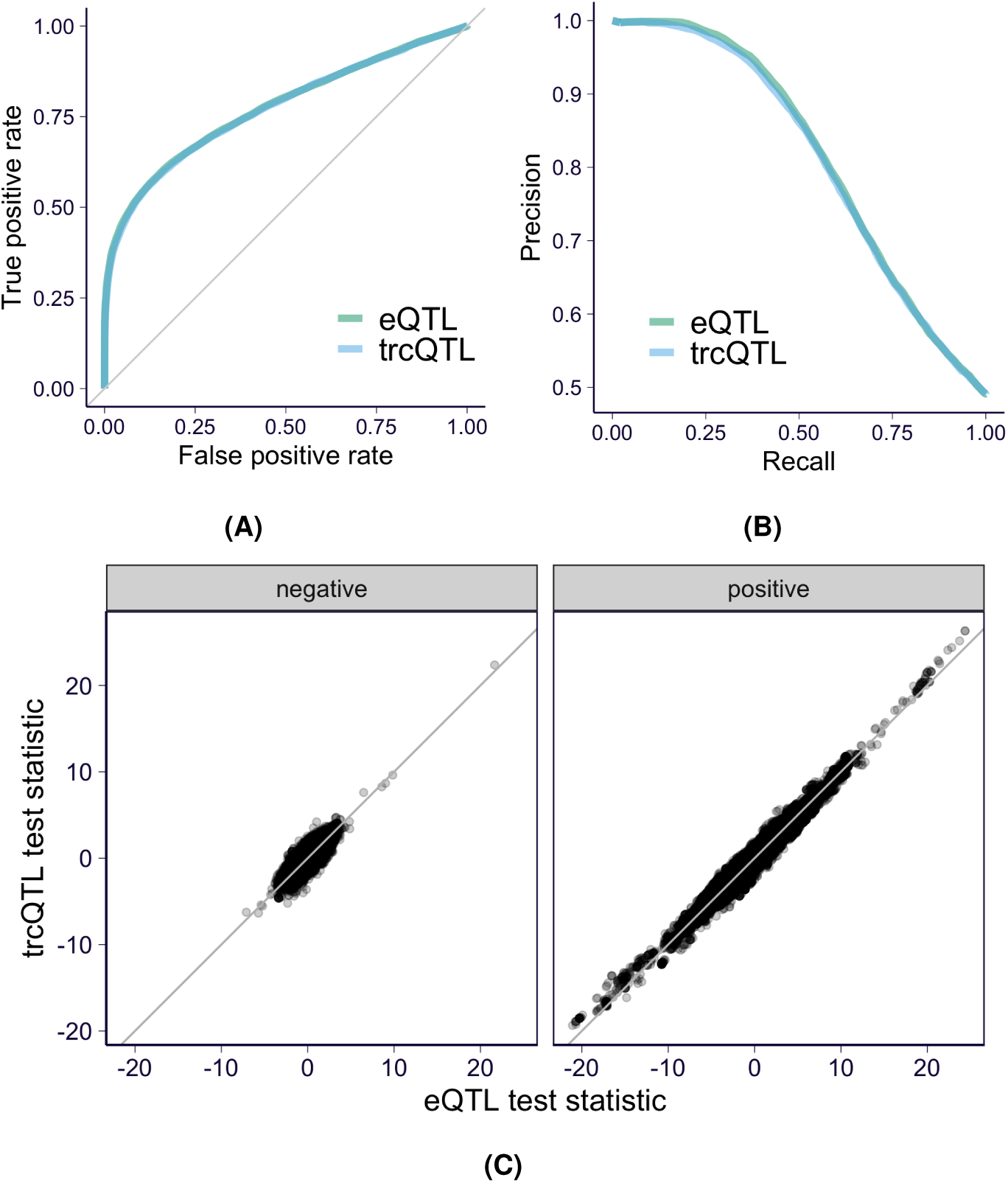
Performance of trcQTL and the standard eQTL approach on genes with low total read counts. Genes with low total counts are defined as having no more than 50 total read counts in any one sample. In GTEx v8 whole blood samples, we extracted 912 genes with low total counts and calculated trcQTL estimates for variants in the corresponding cis-windows. To compare the power of trcQTL and eQTL, we used the 85,129 variant/gene pairs with FDR < 0.05 in eQTLGen as a “ground truth” set. We also randomly selected 88,242 variant/gene pairs from the pairs with p-value > 0.5 in eQTLGen as a negative set. **(A**,**B)** ROC and PR curves for trcQTL and the standard eQTL method. **(C)** Test statistics for the standard eQTL method (x-axis) and trcQTL (y-axis). The variant/gene pairs in the eQTLGen negative set are shown in the left panel, and pairs in the “ground truth” set in the right panel.

**Supplementary Fig. S10.**
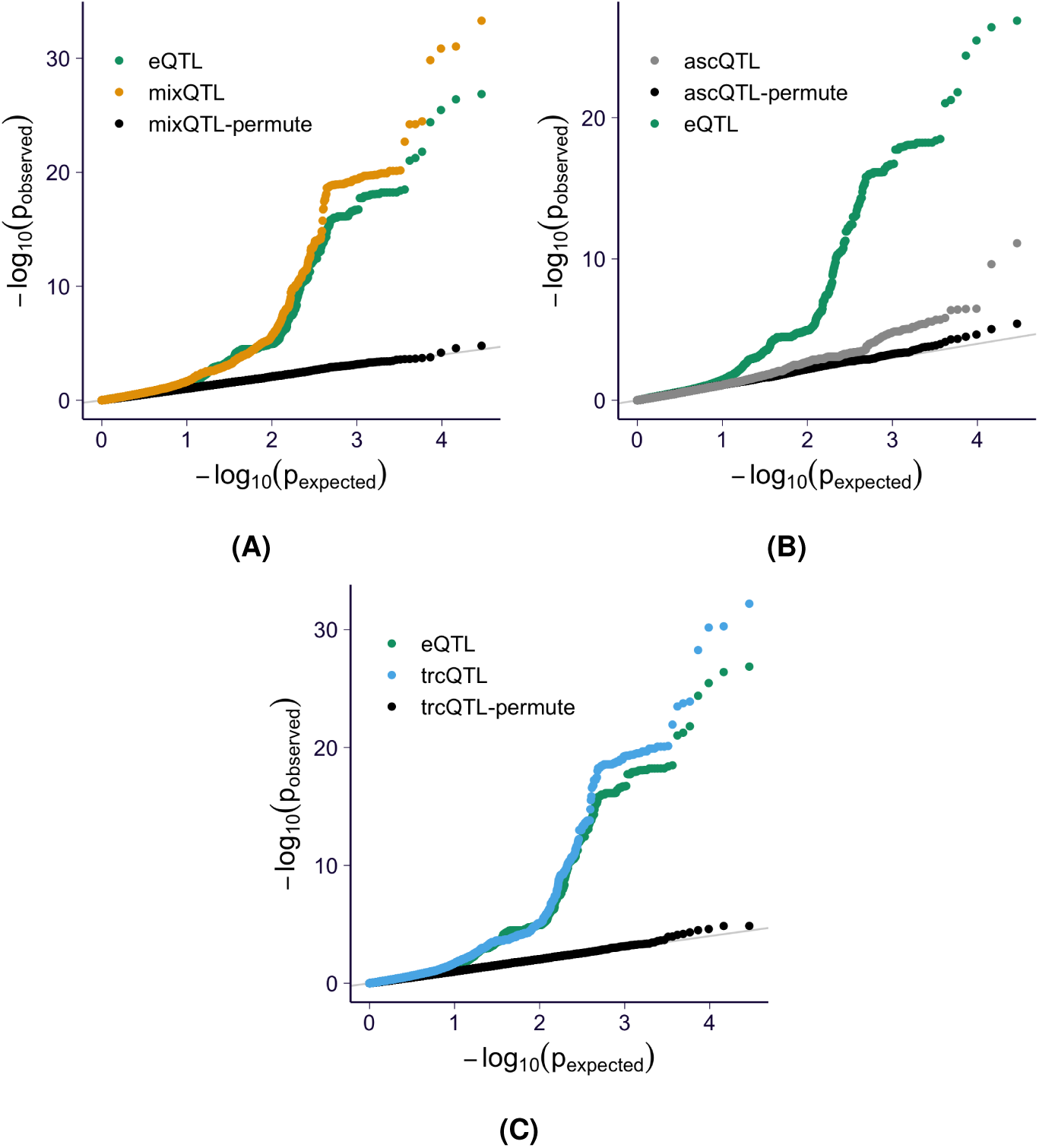
QQ-plot of nominal p-values from ascQTL and trcQTL on four randomly selected genes in GTEx v8 whole blood RNA-seq. The nominal p-values of trcQTL and ascQTL are compared against the standard eQTL method for four randomly selected genes ENSG00000000457, ENSG00000001461, ENSG00000002834, and ENSG00000277734. The results of ascQTL and trcQTL on permuted genotypes are shown in black. **(A)** Results from mixQTL. **(B)** Results from ascQTL. **(C)** Results from trcQTL.

**Supplementary Fig. S11.**
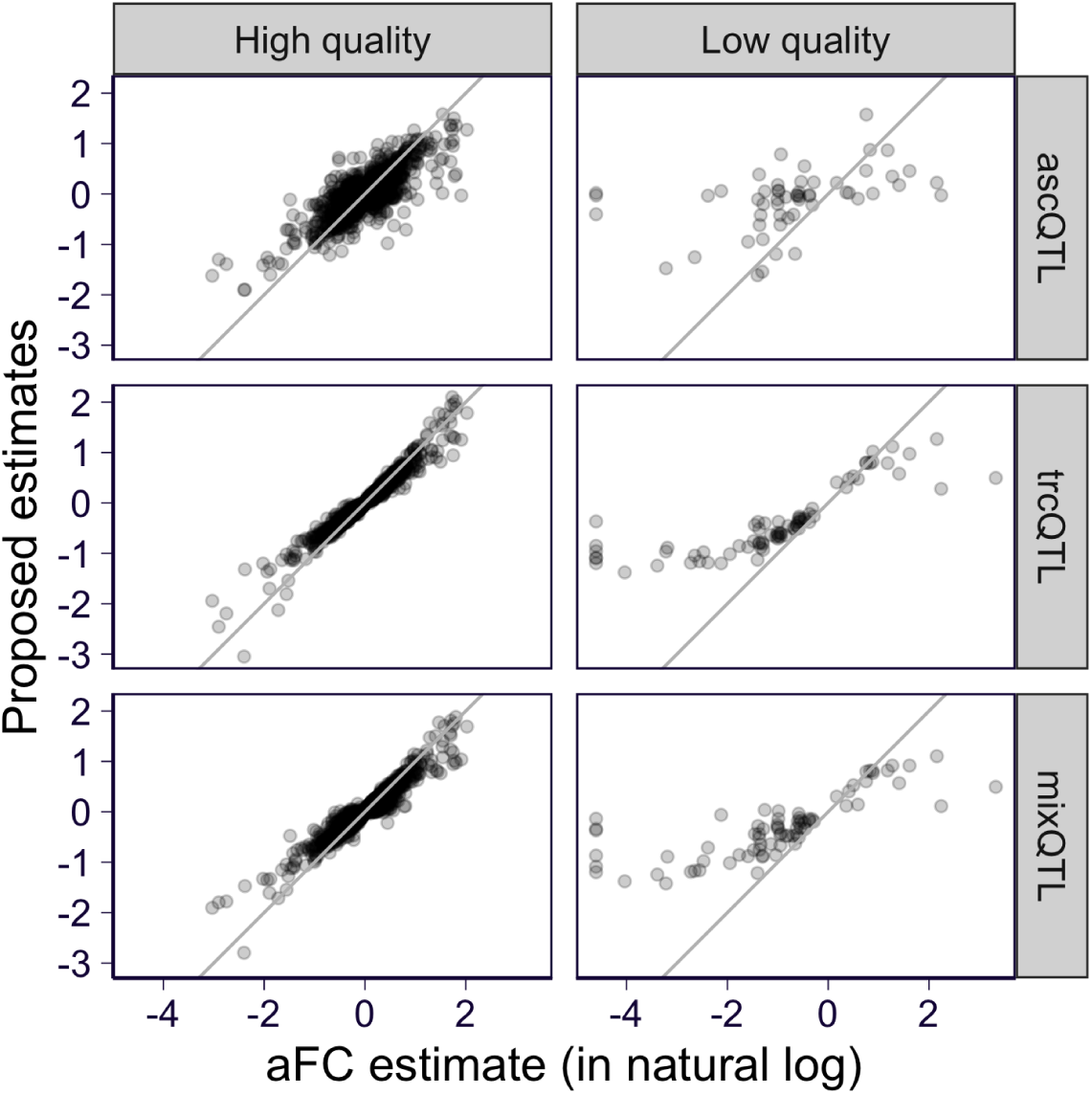
Comparison of aFC estimates from GTEx v8 and the estimated allelic fold change of ascQTL, trcQTL, and mixQTL. The estimates of the top variants in the eGenes of GTEx v8 whole blood are shown (based on eQTL results). On the x-axis, the aFC estimate reported by GTEx v8 is shown (the reported value is in log_2_ and, for visualization, we rescale it to natural log scale by multiplying the value with log(2)). On the y-axis, the estimated allelic fold changes (in natural log scale) of ascQTL, trcQTL, and mixQTL are shown. The variant/gene pairs are stratified on the basis of the quality of aFC estimate, which is defined as ‘high quality’ if the 95% CI of log_2_ aFC is smaller than 1 and the low and high boundaries of the 95% CI are not more extreme than − log_2_(50) and log_2_(50), and as ‘low quality’ otherwise.

**Supplementary Fig. S12.**
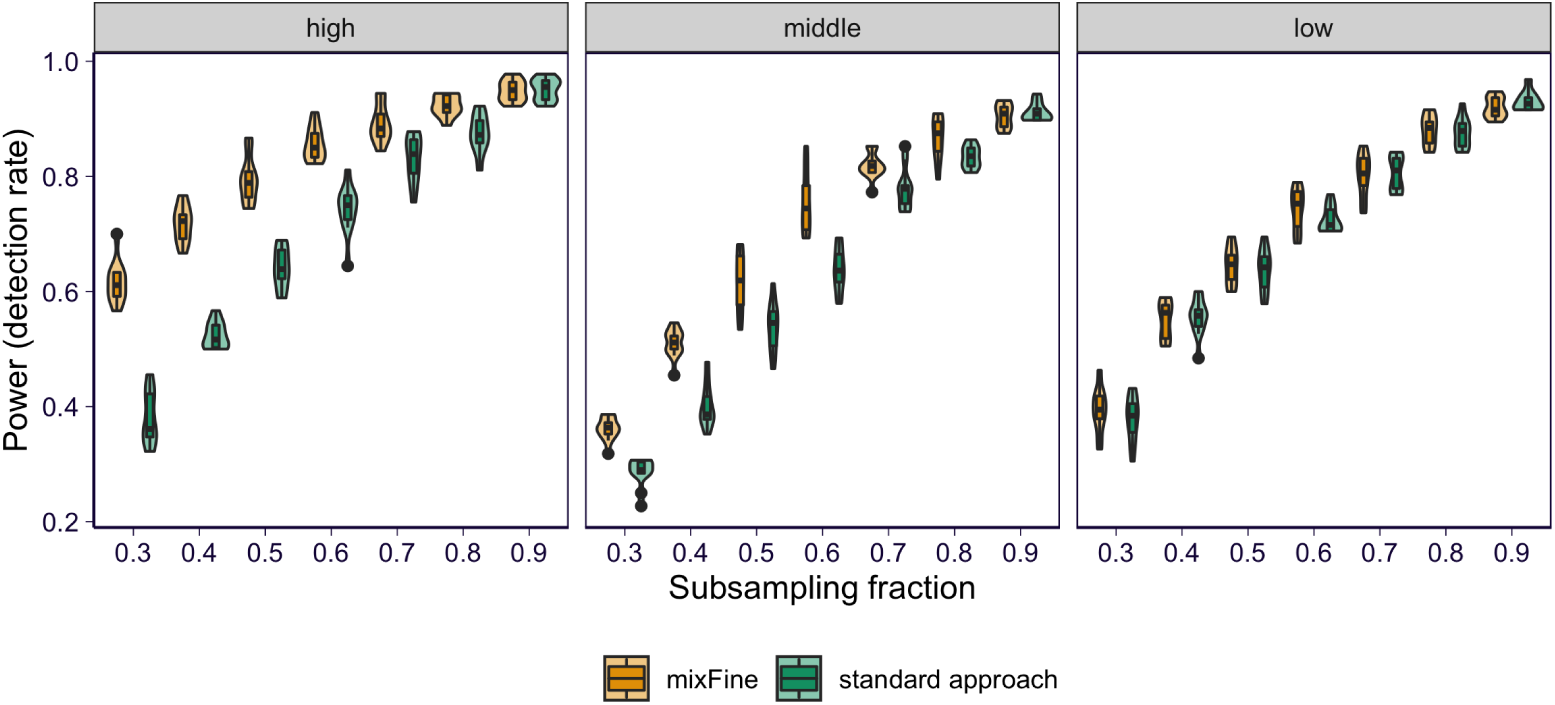
Performance of mixFine on GTEx v8 whole blood RNA-seq stratified by expression level. At each subsampling level (x-axis), the fraction of “consensus SNPs” being detected is shown on the y-axis. Each panel shows the results of genes stratified by expression level tertiles.

**Supplementary Fig. S13.**
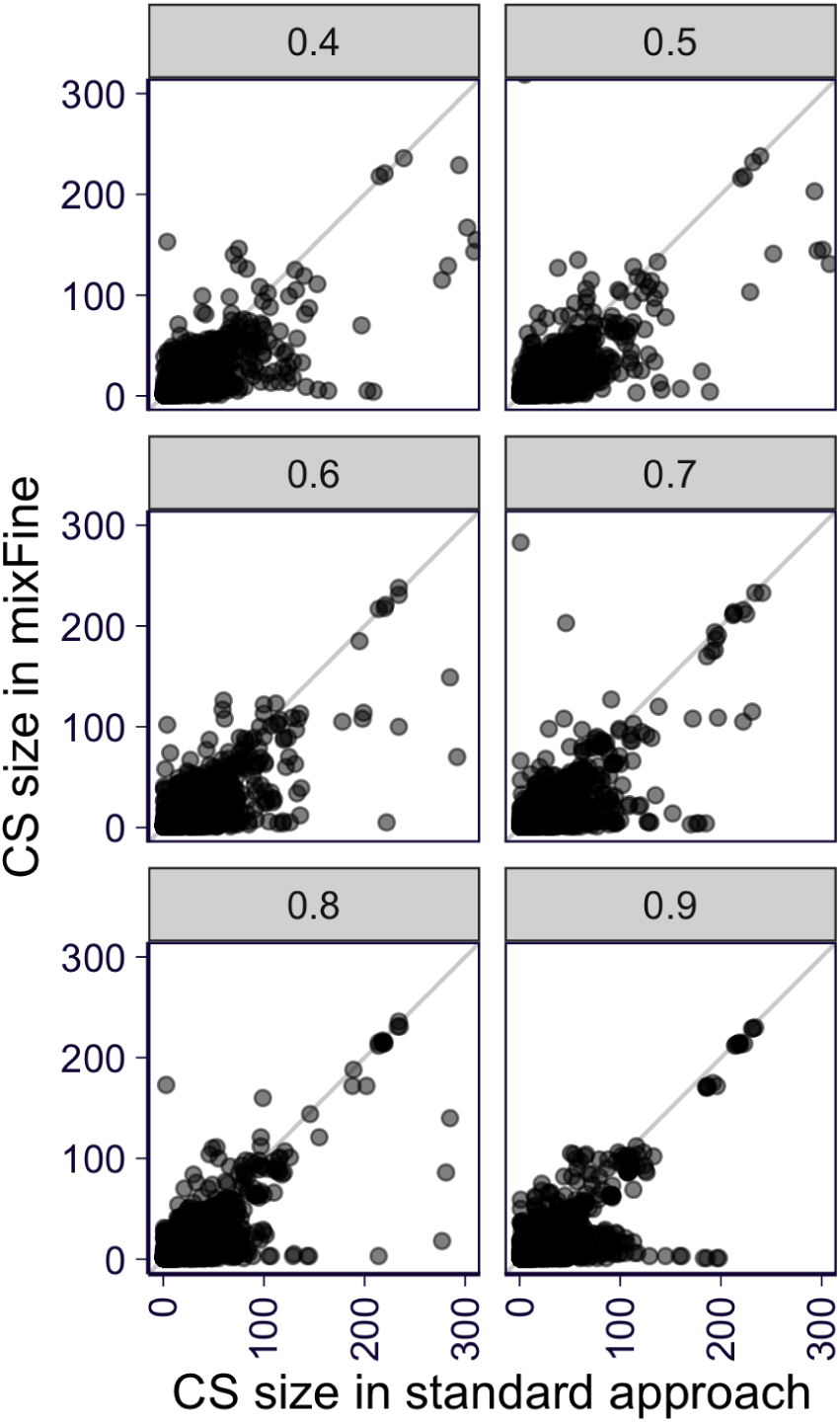
Performance of mixFine on GTEx v8 whole blood RNA-seq on pinpointing the “top” SNPs. At each subsampling level (shown in each panel), we compare mixFine (y-axis) and the standard method (x-axis) on the size of 95% CS’s which are paired by sharing the same “top SNP”.

**Supplementary Fig. S14.**
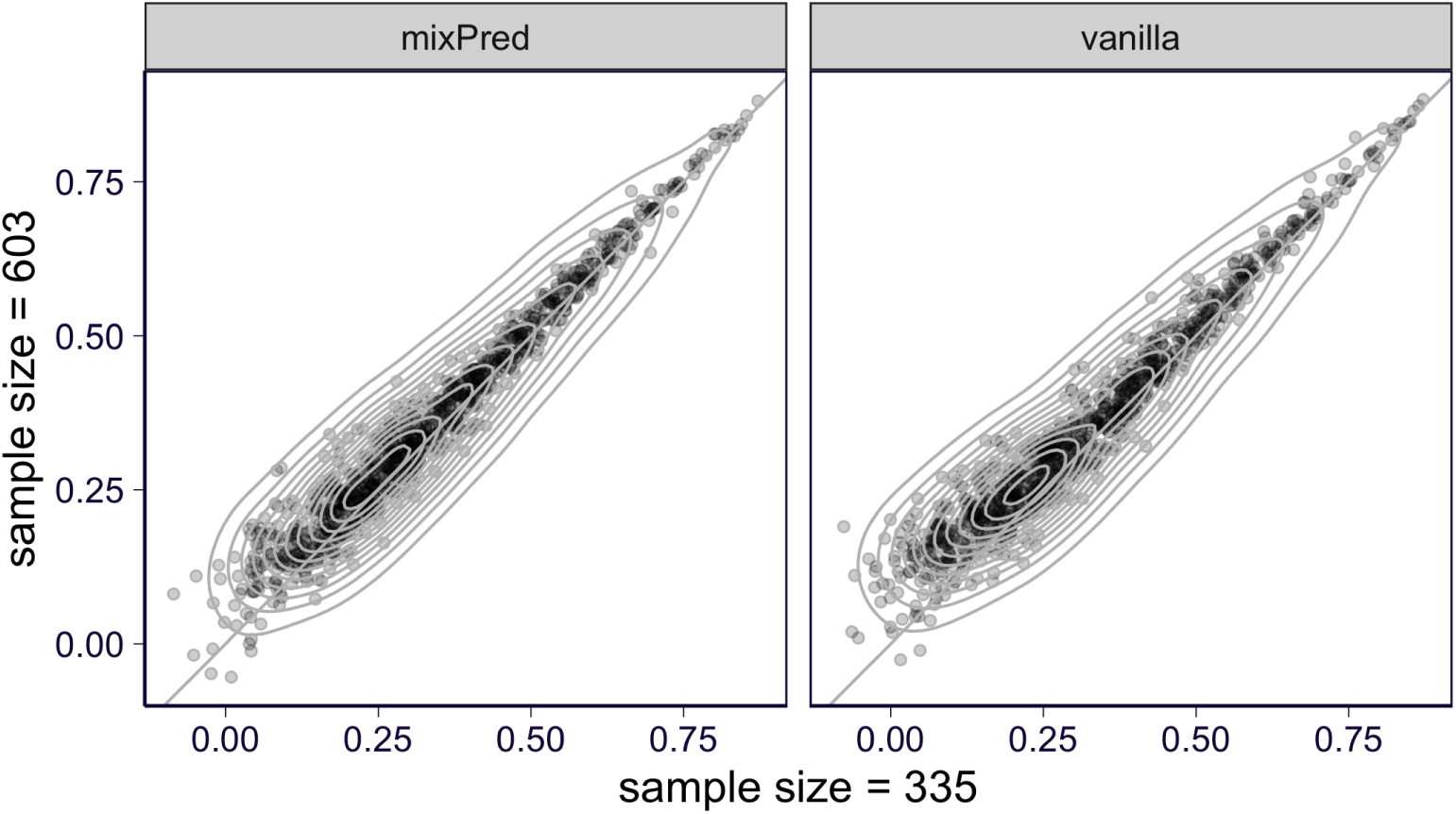
Performance of mixPred and the standard method on GTEx v8 whole blood RNA-seq with different sample sizes. For each gene, the median prediction performance across the *k* held out folds is shown. Here the results with sample size = 335 (*k* = 2) are shown on x-axis and the results with sample size = 603 (*k* = 10) are shown on y-axis. The left panel shows the results of mixPred and the right panel shows the results of the standard method.

## Supplementary Notes

## 7 Statistical model for read count

Here we introduce the statistical model of read count in this paper. For completeness, we opt for keeping some text that overlaps with main text. Recall that *i* indexes individual and *h* indexes haplotypes. 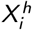 is the phased genotype of the corresponding individual *i* haplotype *h*. 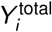 is the total read count within the gene body and *L*_*i*_ is the library size. 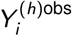 is the allele-specific read count of the corresponding haplotype transcript *h* and 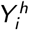 is the actual (though unobserved) read count of the haplotype transcript *h. α*_*i*_ is the expected fraction of allele-specific reads in individual *i*. Additionally, the cis-genetic effect a single SNP on haplotype *h* is represented as 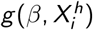 where

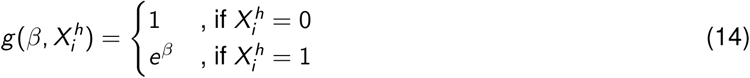

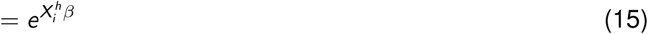

We assume multiplicative effect when there are multiple causal SNPs. And the effect of multiple SNPs *j* = 1, …, *p* is

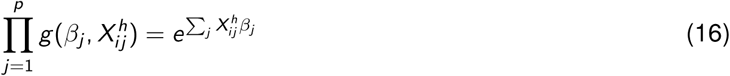

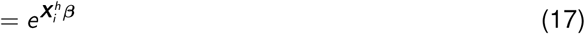

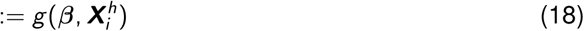

### 7.1 Overview

We model haplotypic count 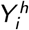 as lognormal distribution as follow.

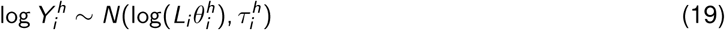

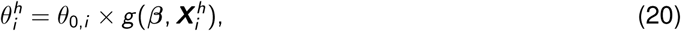

*θ*_0,*i*_ is the baseline abundance of haplotype transcript without considering genetic effect (*i.e*. it represents the abundance when the affecting SNP is reference allele).

In practice, we do not observe 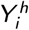 but allele-specific read count 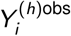. So, we further assume that the baseline abundance of corresponding allele-specific reads are 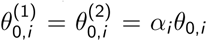. And by definition, total read count 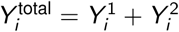. So, similar to Eq 19, 20, 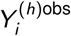 and 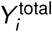 follow

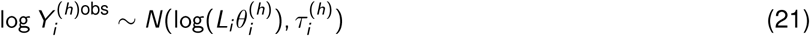

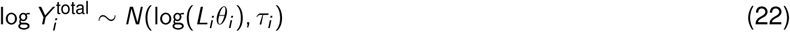

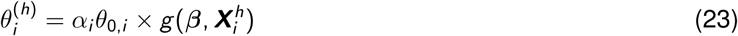

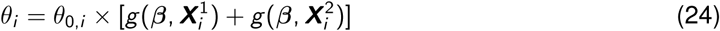

### 7.2 Parameterizing *τ* to weight total and AS count properly

Note that lognormal distribution has the following property.

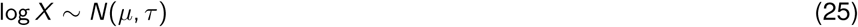

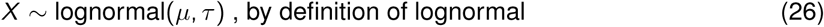

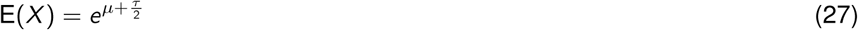

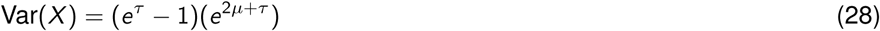

When modeling read count, given the mean, we would like the variance to scale linearly with the mean (as assumed in RASQUAL [Kumasaka et al., 2016]). In other word, we want to ensure that Var(*X*)*/*E(*X*), also known as over-dispersion parameter, is roughly a constant. From Eq 27, 28 we have Var(*X*) = (*e*^*τ*^ −1)E(*X*)^2^. For count data, since *τ* is capturing the variation of count in log-scale, *τ* is typically close to 0. So *e*^*τ*^ − 1 ≈ *τ* and Var(*X*) ≈ *τ* E(*X*)^2^. This result suggests that to ensure Var(*X*)*/*E(*X*) = constant, *τ* should be approximately proportional to 1*/*E(*X*). So, for the distribution of *Y* ∼ lognormal(log(*Lθ*), *τ*), we impose the constraint on *τ* such that *τ* ≈ *σ*^2^*/*E(*Y*). In practice, E(*Y*) is unknown so that we plug-in *Y* in replace of E(*Y*).

## 8 Single-SNP model

On the basis of the model described in Supplementary Notes 7.1, we propose the single-SNP model where we focus on one “test SNP” 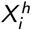 instead of the whole phased haplotype 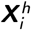. Hence, the cis-genetic effect of interest is 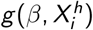.

### 8.1 From likelihood to linear mixed model

Here, we model cis-genetic effect of test SNP as allelic fold change (aFC) [Mohammadi et al., 2017]. So *β* is log-scale aFC in 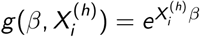. From Eq 21, 23, we have (for *h* = 1, 2)

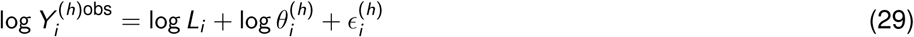

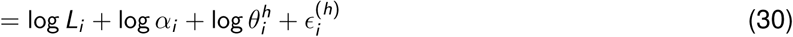

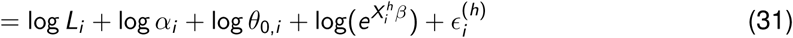

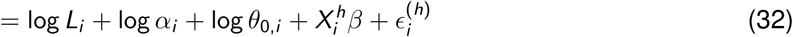

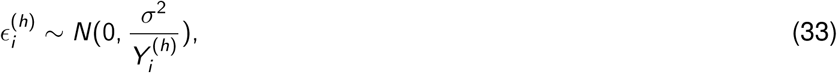

where the error term scaling in Eq 33 follows from the discussion in Supplementary Notes 7.2. To further simplify the term log *θ*_0,*i*_, as the variation of baseline abundance among individuals, we assume 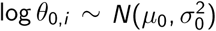. So that Eq 32, 33 can be further written as

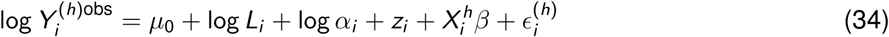

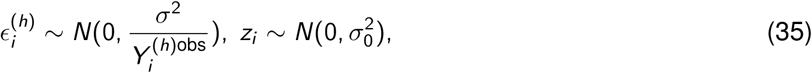

which is the approximated likelihood function for allele-specific counts 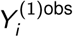 and 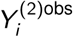. Such likelihood function is equivalent to linear mixed effects model.

Furthermore, we can linearize the likelihood of total read count 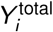 in similar fashion. From Eq 22, 24, we have

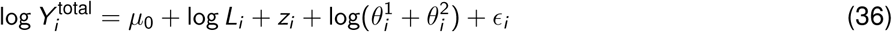

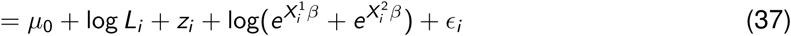

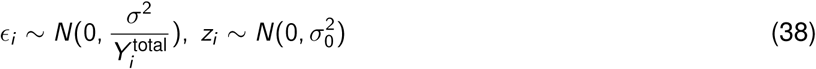

Here we linearize 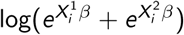 under the weak-effect assumption as follow

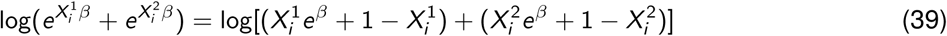

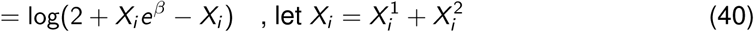

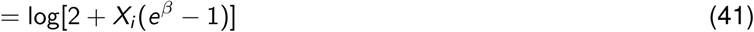

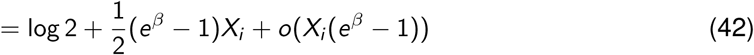

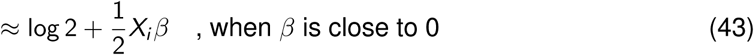

So that Eq 37 can be approximated as

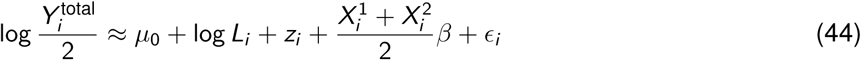

In summary, combining Eq 34,38, 35, 44, we have a linear mixed effects model unifying total and allele-specific read counts after linearization along with other approximations. And it also serves as an approximated likelihood for total and allele-specific reads, in which we can see that these read counts are not independent since they share the same random effect *z*_*i*_.

### 8.2 Simplifying the model

Note that *α*_*i*_ is not observed so that we are unable to solve the model proposed in Supplementary Notes 8.1 in a computationally efficient manner. Here we address this problem by re-parameterizing the model. In principle, conditioning on genetic effect *β*, the ratio of allele-specific reads should be independent to the observations on the total read counts. This intuition motivates us to model the ratio of 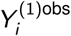 and 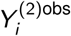 rather than each of them separately. Mathematically, we subtract 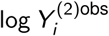 from 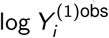, which gives

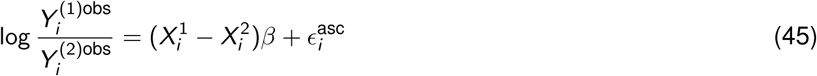

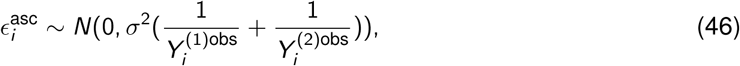

where both *z*_*i*_ and *α*_*i*_ cancel out. This result naturally shows that the likelihood function of 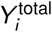 and 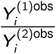 takes the form:

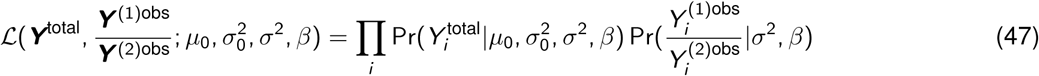

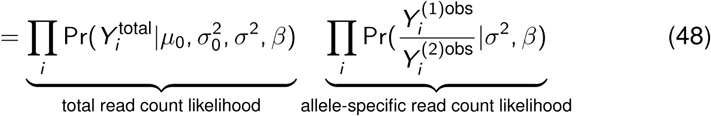

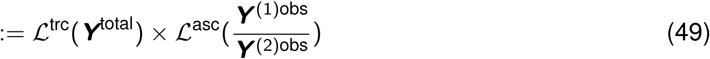

With the simplification shown in Eq 45, the model used for inference can be summarized as follow

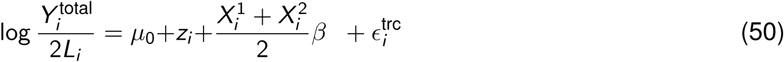

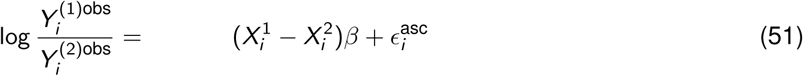

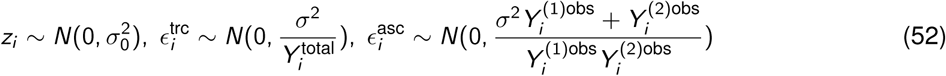

## 9 Generalizing to multi-SNP model

The linearized model described in Eq 50, 51, 52 is easily extensible to multi-SNP scenario since we assume multiplicative genetic effect, as described in Supplementary Notes 18. To see the extension, all we need to examine is how 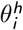 and 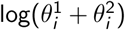 as compared to the single SNP case since the rest of the terms stay the same.

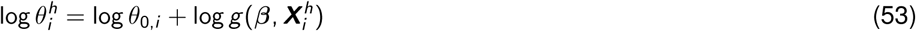

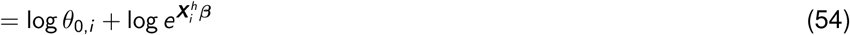

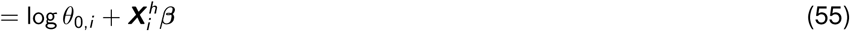

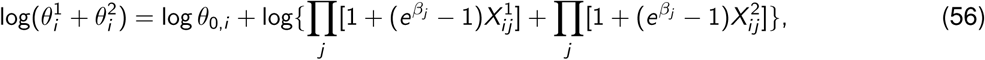

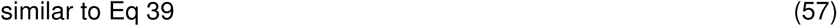

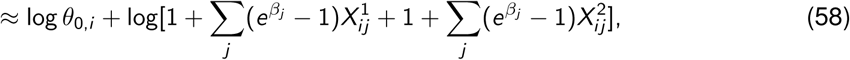

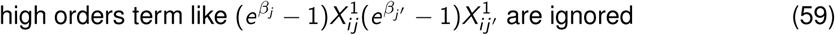

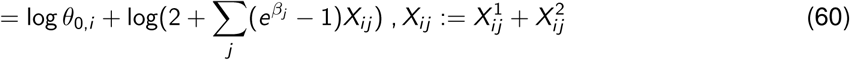

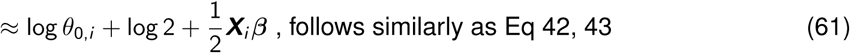

So, we can simply plug-in the multi-SNP version of 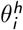 and 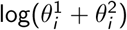 to Eq 30 and 36 respectively and the similar conclusion follows with ***X*** and ***β*** in replace of *X* and *β*.

## 10 QTL mapping procedure

In the following, we describe the mixQTL procedure to map cis-eQTLs under the model proposed in Eq 50, 51, 52.

### 10.1 Converting the problems into two linear regressions

Instead of solving the proposed mixed effects model using numerical solver, we propose a meta-analysis procedure. In this procedure, we solve Eq 50 and 51 separately and meta-analyze the estimates afterwards. Specifically, to solve Eq 50, we first recognize that 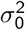 is much larger than 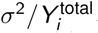. This is due to the following three facts: 1) *σ*^2^ has the scale of 1 (*σ*^2^ = 1 corresponds to Poisson); 2) 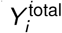 is total count which is typically hundreds to thousands; and 3) 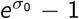 is roughly the scale of the ratio between *θ*_0,*i*_ and the population mean abundance (E(*θ*_0,*i*_)), which makes *σ*_0_ ∼ 0.5 (corresponds to *θ*_0,*i*_ to E(*θ*_0,*i*_) ratio being from 0.6 to 1.6) a reasonable estimate. So, we further simplify Eq 50 by ignoring the noise term from 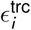. Such simplification results in the following linear model

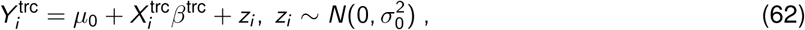

where 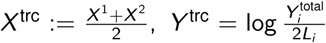. Eq 62 itself can be used for QTL mapping and we call this approach trcQTL in the paper.

For solving Eq 51, notice that it is weighted simple linear regression with the form

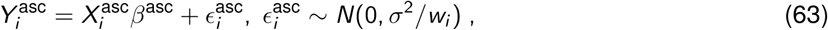

where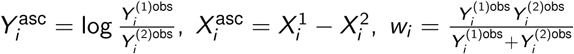. We call QTL mapped by Eq 63 ascQTL.

Note that we can combine Eq 62 and 63 and solve them jointly in close form. But here we still prefer meta-analysis for two reasons: 1) it allows combining summary statistics across studies; and 2) it allows the over-dispersion in total and allele-specific read counts to be different which is more realistic in practice since total and allele-specific read counts may go through different pre-processing steps.

Since the inference of linear regression has analytical solution which only involves *X*^*T*^ *X* and *X*^*T*^ *Y*, we can solve it quickly and in a parallel way as proposed by Matrix eQTL [Shabalin, 2012]. We sketch the pseudocode on calculating trcQTL and ascQTL estimates in matrix form in Supplementary Notes 13.

### 10.2 Meta-analysis for QTL mapping

Once we obtain estimated 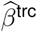 and 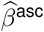, we can use these estimates to approximate ℒ^trc^ and ℒ^asc^ in Eq 49. Specifically, when sample size is large,

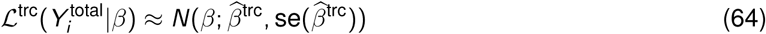

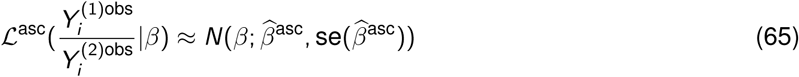

So that the joint likelihood, as factorized in Eq 48, is simply 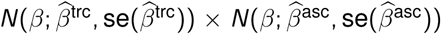. As shown previously [Lee et al., 2016], maximizing the approximate joint likelihood is equivalent to inverse-variance meta-analysis, which takes the form

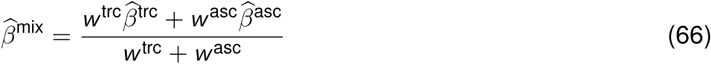

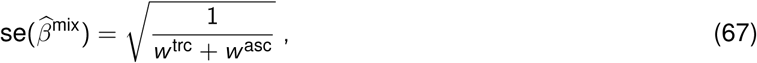

Where 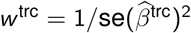 and 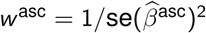.

## 11 Inference procedure for multi-SNP model

With the simplification made in Supplementary Notes 10.1, the multi-SNP model can be written as

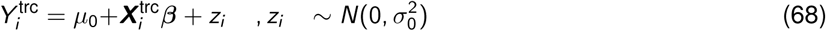

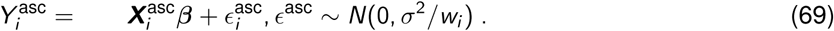

### 11.1 Motivating two-step inference procedure

Here we focus on two inference problems under the multi-SNP model: 1) construct genetic predictor of expression; and 2) infer whether *β*_*k*_ is non-zero, *i.e*. causal SNP. Problem 1) is prediction problem in machine learning context and in terms of building genetic predictor, elastic net has been used for this task [Gamazon et al., 2015]. For problem 2), the inference problem is formulated into a Bayesian variable selection problem and efficient solvers such as susieR [Wang et al., 2019] and DAP-G [Lee et al., 2018] have been developed in the context of eQTL analysis.

However, the existing methods only use total read information (typically inverse normalized expression) and they assume the inversely normalized expression *Y* and genotype vector ***X*** follow *Y* ∼ *N*(***Xβ***, *v*). The modeling assumption is very close to Eq 68, 69 but it requires equal variance in error term and shared intercept across all observations. To apply the existing tools, we need to bypass the gap between our model and their modeling assumption. For this reason, we propose a two-step inference procedure to perform inference for multi-SNP model. In step 1, we infer *μ*_0_ 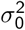, and *σ*^2^ and transform the data such that they approximately follow *Y* ∼ *N*(***Xβ***, *v*). And in step 2, we apply the transformed data to existing solvers for both prediction and fine-mapping problems.

### 11.2 Inferring 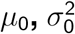, and *σ*^2^

To estimate *σ*^2^ from Eq 69 is equivalent to estimate the mean squared error (MSE) of the model. To avoid overfitting ***β*** and underestimating MSE, we apply 4-fold cross-validation using LASSO to get effect size estimate 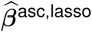. And 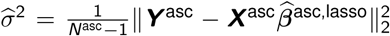 where *N*^asc^ is the number of allelic imbalance observations. Any alternative approach would be to treat 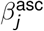 as random effect and estimate *σ*^2^ as one of the variance components. But here we implement the former.

We apply similar approach to estimate 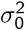 using LASSO. To obtain *µ*_0_, we fit Bayesian variable selection model using susieR with the total read count observations 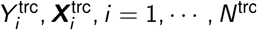. The output intercept 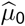 is the estimate.

### 11.3 Data transformation and inference

Once we obtain 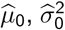, and 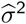, we shift and re-scale the total and allelic imbalance observations by

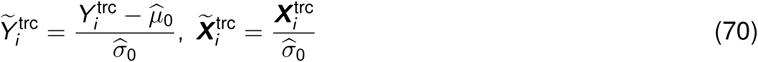

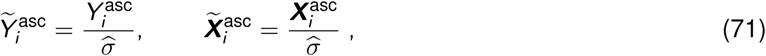

where the transformed data (on the left-hand side) is used for downstream analysis on performing prediction and fine-mapping.

Specifically, we concatenate 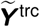 and 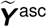 into one vector 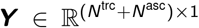 and similarly we concatenate 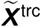 and 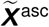 into one matrix 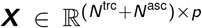 where *p* is the number of SNPs. To perform fine-mapping, we run `susieR::susie(X = X, Y = Y, intercept = FALSE, standardize = FALSE)` with `X` equal to ***X*** and `Y` equal to ***Y***. To build prediction model, we run `glmnet::glmnet(x = X, y = Y, lambda = lambda, alpha = 0.5)` with `x` equal to ***X*** and `y` equal to ***Y***. The hyperparamter `lambda` is selected by 5-fold nested cross-validation where at each `lambda` the 5-fold cross-validation are repeated three times and `lambda` that has lowest cross-validated mean squared error (averaged across three runs) is used. For comparison, we feed the part of total read count data (***X***^trc^, ***Y*** ^trc^) directly into: 1) susieR for fine-mapping; and 2) elastic net for prediction. The procedure is the same but ***X, Y*** are replaced by ***X***^trc^, ***Y*** ^trc^. And we call this total read count-only approach for fine-mapping and prediction as trcFine and trcPred.

## 12 Simulating RNA-seq reads

To examine the performance of the methods, we propose and implement a simulation scheme which generates total and allele-specific read counts. The simulation procedure includes three parts: 1) simulate gene body which will be aligned by reads; 2) randomly draw the causal variants; 3) simulate the number of reads for each haplotype transcript and place these reads to the gene body obtained in step 1). The total and allele-specific read counts can be directly read out from step 3) where the total read count is the sum of two haplotypic read counts and the allele-specific read count is the number of reads overlapping with heterozygous sites within gene body.

In step 1), we fix the length of gene body to be 10kbp. To simulate the heterozygous sites within gene body for each individual, we start with determining the position of polymorphic sites along gene body. We first sample the number of polymorphic sites from Binomial distribution, and then draw their positions and minor allele frequencies (MAFs). And finally, whether a polymorphic site is heterozygous in an individual is determined by Bernoulli distribution with MAF. The procedure is sketched as follow.

1. Number of polymorphic site within gene body *N*_*h*_ ∼ Binomial(*L*_gene_, *f* ^*h*^), where *L*_gene_ = 10^4^, *f* ^*h*^ = 0.001.
2. Position *P*_*m*_ (*m* = 1, …, *N*_*h*_) of these polymorphic sites are sampled by *P*_*m*_ ∼ Sample({1, …, *L*_gene_})And the corresponding MAF *f*_*m*_ are drawn from *f*_*m*_ ∼ Uniform(maf^*l*^, maf^*h*^), where maf^*l*^ = 0.05, maf^*h*^ = 0.3.
3. For each individual *i*, whether the *m*th polymorphic site is heterozygous (denote as *Z*_*im*_) is determined by *Z*_*im*_ ∼ Bernoulli(2*f*_*m*_(1 − *f*_*m*_)).

In step 2), the genetic effect equals to 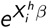 (in single-SNP model) and 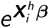 (in multi-SNP model). To do so, we need to obtain haplotype and effect size. For single-SNP model, we first sample MAF of the causal variants and obtain the two haplotypes of each individual by drawing from Bernoulli. For multi-SNP model, we use the 1000G phase3 genotypes of European individuals. In brief, we randomly select 200 genes on chromosome 22 and extract phased genotypes of 1Mbp cis-window surrounding the transcription start site of them (excluding variants with allele frequency < 0.01 or > 0.99). The genetic effect size, *e*^*β*^, ranges among 1, 1.01, 1.05, 1.1, 1.25, 1.5, 2, 3 for single-SNP case. In multi-SNP case, the number of causal SNPs is sampled from 1, 2, 3 and the genetic effect ranges from 0.015 to 0.075 such that the heritability ranges approximately from 19.4% to 54.5%. The detailed procedure for sampling 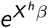 and 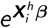 is as follow.

- **Single-SNP scenario**:
  1. Sampling 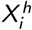: MAF of causal SNP *f* ^*c*^ ∼ Uniform(maf^*l*^, maf^*h*^) and 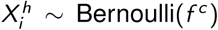 Bernoulli(*f* ^*c*^) where maf^*l*^ = 0.05, maf^*h*^ = 0.3.
  2. Setting up *β*: fixed to 1, 1.01, …, 2, 3.
- **Multi-SNP scenario**:
  1. Sampling 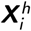 : obtained from 1000G phased genotypes.
  2. Setting up ***β***: number of causal SNPs ∼ Sample({1, 2, 3}) and the genetic variation *v*_*g*_ ∼ Uniform(0.015, 0.075). The genetic effect of causal variants are determined by randomly partition the genetic variation and convert per-SNP genetic variation into effect size by 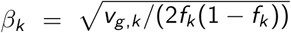 where *f*_*k*_ is MAF of *k*th causal SNP.

In the step 3), the last step, we sample the reads coming from each of the haplotype transcripts. The procedure is as follow.

1. For individual *i*, sample library size *L*_*i*_ ∼ NegativeBinomial(size, prob) where size = 15, prob = 1.6 × 10^−7^ (Negative Binomial follows parameterization in rnbinom in R).
2. And then, sample individual-specific baseline abundance *θ*_0,*i*_ ∼ Beta where E(*θ*_0,*i*_) ranges among 5 × 10^−5^, 2.5 × 10^−5^, 1 × 10^−5^, 5 × 10^−6^, 2.5 × 10^−6^, 1 × 10^−6^ and sd(*θ*_0,*i*_) = E(*θ*_0,*i*_)*/*4 (so that the non-genetic variation is roughly 1*/*4^2^ = 1*/*16).
3. The actual relative abundance of haplotype *h* in individual *i* is 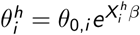 or 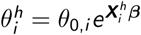
4. Sample actual read count for each haplotype: 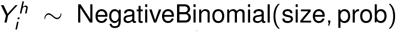 where size = 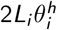, prob =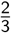. This corresponds to 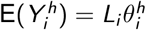 and Var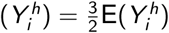.
5. Randomly place reads, 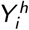 in total, onto the corresponding gene body simulated in step 1) where the read is aligned to each position of gene body with equal probability.
6. Total count is 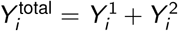 and allele-specific count 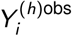 is the number of reads (as part of 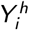) that overlaps with the heterozygous sites of the individual (indicated by *Z*_*i·*_).

## 13 Pseudocode on solving trcQTL and ascQTL in matrix form

We sketch the matrix operations for solving a grid of least squares problems ***y***_*k*_ ∼ ***x***_*j*_ for each pair of *j, k* where we let *Y* = [***y***_1_, …, ***y***_*K*_] and *X* = [***x***_1_, …, ***x***_*n*_]. To obtain nominal p-value, *K* = 1. For permutation procedure proposed in fastQTL [Ongen et al., 2015], *K* equals to the number of permutation and ***y***_*k*_ is the *k*th permuted ***y***.

To ensure trcQTL and ascQTL ran on the same permuted ***y***, we perform permutation before removing low count observations. So that in each permutation, different individuals are removed by low-count filter. To account for this fact, we introduce mask *M* ∈ {0, 1} ^*n*×*K*^ where *M*_*ik*_ indicating if the *i* th individual is included in *k*th permutation.

For trcQTL, the corresponding least squares problem has intercept, as mentioned in Eq 62. The pseudocode to solve the grid of trcQTL problems for all cis-SNP of a gene is sketched in Algorithm 1 where *Y* = ***Y*** ^trc^ for nominal pass and *Y*_*·k*_ = *P*_*k*_ ***Y*** ^trc^ with permutation matrix *P*_*k*_ for permutation pass.

Note that the pseudocode only requires basic matrix operation. The matrix operation is elementwise if not notice explicitly. The Einstein summation is represented by `einsum` with similar arguments as `numpy.einsum` in Python. For instance, `einsum(‘ij,jk → ik’, A, B)` means that to take the inner product of the *i* row in A and *k* column in B as the element at *i* th row and *j*th column in the output matrix.

Similar to trcQTL, the corresponding least squares problem of ascQTL is weighted without intercept, as mentioned in Eq 63. The pseudocode to solve the grid of ascQTL problems for all cis-SNP of a gene is sketched in Algorithm 2 where *Y* = ***Y*** ^asc^ for nominal pass and *Y*_*·k*_ = *P*_*k*_ ***Y*** ^asc^ with permutation matrix *P*_*k*_ for permutation pass. And *W* as the weight matrix should be permutate accordingly, *i.e. W*_*·k*_ = *P*_*k*_ ***w***. And to obtain valid mixQTL estimates under permutation, *P*_*k*_ is required to be shared by trcQTL and ascQTL in permutation pass.

Note that both Algorithm 1 and Algorithm 2 are iteration free. And throughout the computation, only two-way tensors are involved explicitly so that the memory usage does not blow up.

### Algorithm 2: Solve multiple least squares problems *y* = *bx* + *e* with weight *w* in matrix form

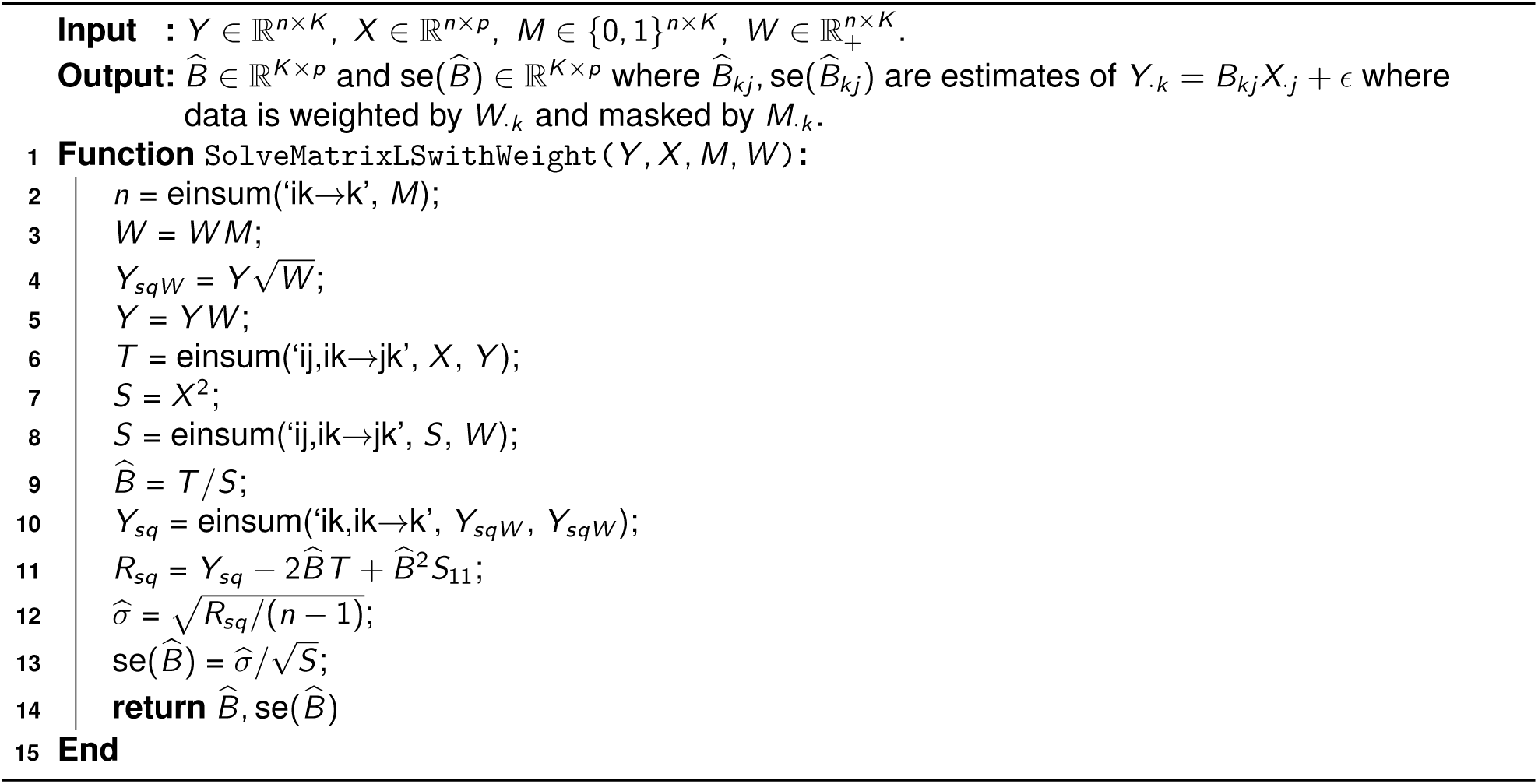

## 14 Evaluating QTL mapping performance using eQTLGen results

To evaluate the performance of QTL mapping method, we treat eQTLGen [Võsa et al., 2018] as a silver standard, in the sense that eQTLs identified as positive in eQTLGen are treated as the true associations and the non-significant variant/gene pairs in eQTLGen are treated as true non-associations. Although 336 GTEx samples are included in eQTLGen analysis, they make up of only around 1.5% of total samples. So, eQTLGen results are unlikely driven by GTEx samples. And besides, GTEx v8 includes additional samples that are not included in eQTLGen. Therefore, eQTLGen is an approximately independent eQTL study with much larger sample size (50-fold relative to GTEx v8) and diverse populations (predominantly Europeans along with other populations).

### Algorithm 1: Solve multiple least squares problems *y* = *a* + *bx* + *e* in matrix form

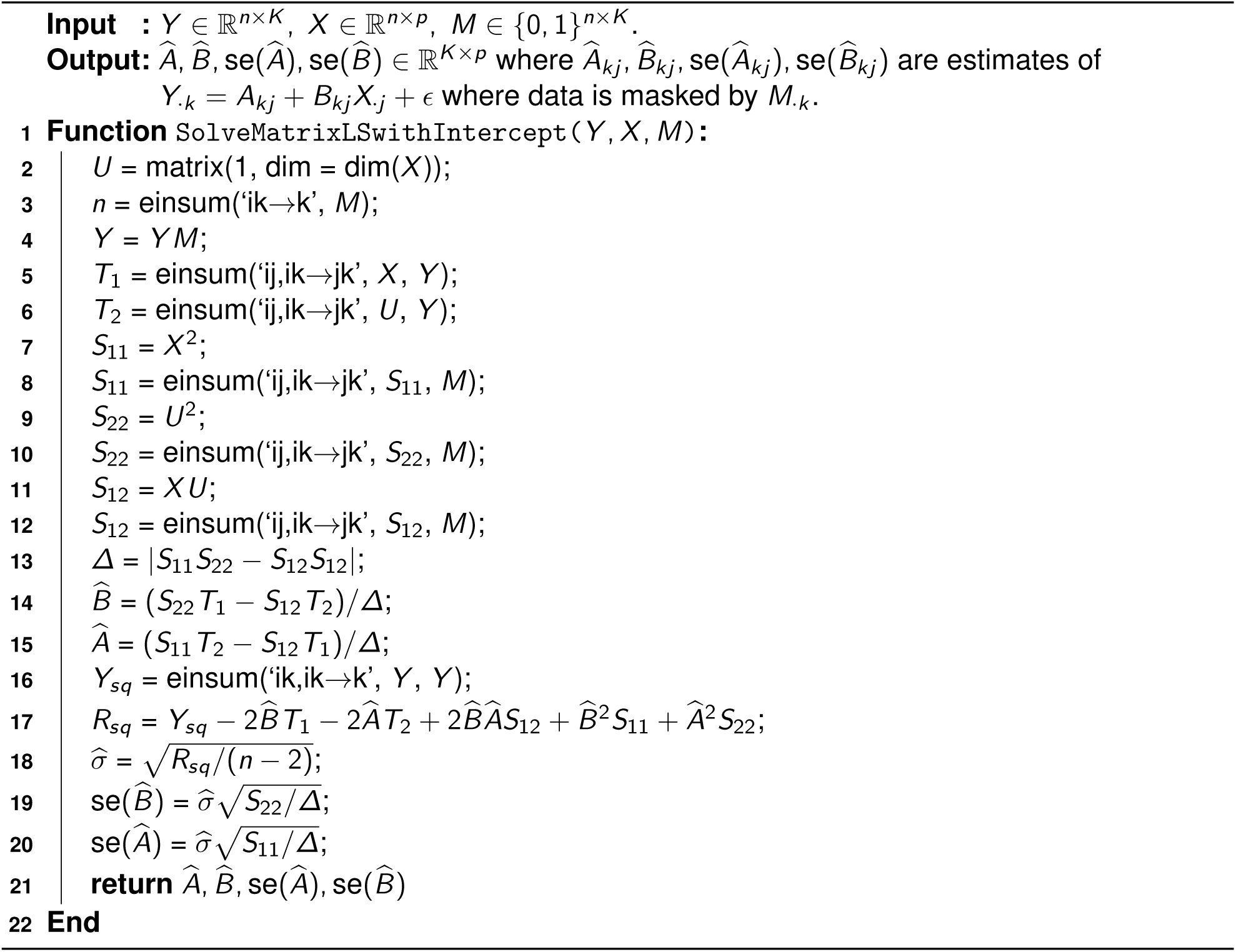

To simplify the analysis, we randomly select 100,000 eQTLGen cis-eQTLs (FDR ¡ 0.05) as the true associations in the silver standard. And we randomly collect 100,000 variant/gene pairs in eQTLGen with p-value ¿ 0.5 as the true non-associations. Among those variant/gene pairs in silver standard, 96,660 true associations and 78,691 true non-associations are included in both our mixQTL mapping pipeline and GTEx v8 analysis. So that we keep only these variant/gene pairs for downstream analysis.

### 14.1 Comparing the effective sample size

To compare the effective sample size between mixQTL and eQTL approaches, we performed analysis similar to [Loh et al., 2018]. Here, we utilize the fact that *χ*^2^ statistic scales proportionally with the sample size, among those true associations. So, we can calculate the ratio 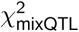 over 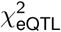 for each truly associated variant/gene pair as the measure of effective sample size of mixQTL relative to eQTL approach. Specifically, we calculate the relative effective sample size using the true associations in the silver standard constructed above (as the proxy of true associations based on independent evidence). Note that the gain of power in mixQTL depends on the amount of allele-specific observations so we measured the average relative effective sample size as the median of the *χ*^2^ ratio. Among the 96,660 variant/gene pairs collected as true associations in silver standard, we measured the median of 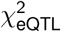 as 2.59 and the median of 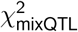 as 3.56. And the median of the ratio 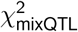 over 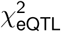 is 1.29. In other word, it suggests that the mixQTL approach (with 670 individuals) is equivalent to the eQTL approach with 863 individuals.

### 14.2 Drawing receiver operating characteristic and precision-recall curves

The ROC and PR curves are constructed using − log(*p*) as prediction score (higher means more likely to be causal). To simplify the calculation, we evaluate the performance measures at a grid of score cutoffs: 0.1, 0.2, …, 1.9, 2, 2.2, …, 2.8, 3, 4, …, 50. For ROC curve, we calculate true positive rate and false positive rate at these cutoffs. And similarly, for PR curve, we calculate precision and power at these cutoffs.

